# What happens when Arctic sea-ice melts? Macroalgae as a summer refuge for sympagic nematode *Theristus melnikovi*

**DOI:** 10.64898/2026.10.16.759596

**Authors:** Vanessa Pitusi, Janne E. Søreide, Rebecca J. Duncan, Oleksandr Holovachov

## Abstract

Sea ice-associated (sympagic) nematodes are particularly vulnerable to ongoing Arctic sea-ice loss, yet the capacity of these sympagic species to persist in ice-free habitats remains poorly understood. We investigated the summer habitat and taxonomic identity of the sympagic nematode *Theristus melnikovi*, in Svalbard, using an integrative taxonomic approach. We barcoded 17 morphologically identified nematodes from macroalgae (5), sea-ice (9), and sediment (3). Morphological analysis and phylogenetic reconstruction based on 18S rRNA gene analyses revealed that the macroalgae-associated *T. melnikovi* represented a habitat-associated phenotypic variation of *T. melnikovi* rather than a morphotype of the benthic nematode *T. acer*. Macroalgae-associated specimens were generally shorter and stouter, with proportionally longer tails and shorter anterior structures than their ice-associated counterparts, resulting in a superficial resemblance to *T. acer.* Despite these morphological differences, molecular analyses consistently placed macroalgae-associated individuals within the *T. melnikovi* clade and haplotype. Our results highlight that *T. melnikovi* may have a greater ecological flexibility than previously assumed and macroalgae may offer an alternative (summer) habitat, or refuge, during ice-free periods. This capacity could become increasingly important as Arctic sea-ice duration and extent continue to decline.

## Introduction

Nematoda represents one of the most taxonomically and functionally diverse phyla, prevalent in communities in freshwater (Traunspurger et al., 2020), marine environments such as beaches (Maria et al., 2016), the deep-sea (Leduc & Rowden, 2018), and sea-ice (Blome & Riemann, 1999; Bluhm et al., 2018). In sea-ice, nematodes live within the intricate network of microscopic brine channels that are created during sea-ice freeze-up. They share this space with fungi, bacteria, viruses, protozoa, protists, and other meiofaunal groups (Lund-Hansen et al., 2024). This community is generally referred to as sympagic (“ice-associated”) and occurs in both offshore and coastal Arctic sea-ice (Bluhm et al., 2018). In the Arctic Ocean, sympagic communities inhabit drift ice that is not attached to land, and landfast ice, which is attached to the shoreline and typically undergoes an annual cycle of winter formation and spring melt. Consequently, sympagic communities in landfast ice are adapted to deal with this seasonality in ice presence and absence in terms of their life history and ecology. These adaptations are becoming increasingly important with ongoing reduction in sea-ice in the central Arctic (Heuzé & Jahn, 2024) and in coastal areas (e.g. Urbański & Litwicka, 2022), as the growth-and-melt cycle of landfast ice has already shifted with later onset of freeze and an earlier melt in many areas of the Arctic (e.g. Mahoney & Einhorn, 2026; Papin et al., 2026). Thus, it is imperative to understand how resilient sympagic nematodes are to their changing ice-habitat, and if they can find refuge in another habitat.

The sympagic nematode *Theristus melnikovi* Tchesunov, 1986 was the first species to be described from perennial sea-ice over the Central Arctic Ocean (Tchesunov, 1986), and since then it has been recorded in multi-year ice from the Fram Strait (Friedrich, 1997), first-year ice in the Barents Sea (Timchenko et al., 2021), seasonal ice from the Laptev Sea (Tchesunov & Riemann, 1995), Resolute Bay in the Canadian Arctic (Riemann & Sime-Ngando, 1997), the White Sea (Tchesunov & Portnova, 2005), the Kara Sea and the East Siberian Sea (Timchenko et al., 2024b), Svalbard (Pitusi et al., 2021), and sediment from the East Siberian Sea (Timchenko et al., 2024b) and White Sea (Timchenko et al., 2025). Thus, this species can be considered to have a pan-Arctic distribution across multiple ice types in which it has been proposed to reproduce based on the presence of males, females, and juveniles. However, its life history is based on scattered observations. In perennial ice, it is assumed that all life stages spend their whole life cycle in the ice (Tchesunov, 1986), whereas the fate of those found in seasonal landfast ice is more uncertain. It has been proposed that after the ice melt, *T. melnikovi* juveniles spend their summer on the seafloor before being incorporated back into sea-ice in the winter (Timchenko et al., 2024b). However, this is speculative and we have no concrete data to confirm this theory, and the abundance of *T. melnikovi* found in sediment is very low (Timchenko et al., 2024b; Timchenko et al., 2025). Similarly to its life history, there is little information on its feeding ecology. It is assumed to absorb dissolved organic matter, feed on bacteria and ice algae based on gut content analysis and resource availability (Tchesunov & Riemann, 1995; Kramer, 2011).

These uncertainties concerning its life history and ecology make it difficult to assess how *T. melnikovi* may respond to changing sea-ice conditions in the Arctic. The low abundances in sediment may point towards another summer habitat that has not been explored yet. We know from the literature that a suitable alternative benthic habitat for nematodes is macroalgae, because they are structurally complex (Heip et al., 1985). Consequently, macroalgae not only offer refuge from predators, but also offer a variety of food resources such as bacteria on biofilms, detrital material or other metazoan inhabitants (Heip et al., 1985). *Theristus* Bastian, 1865, and more specifically, the species *Theristus acer* Bastian, 1865 has been documented from a variety of intertidal habitats around the globe including macroalgae (Kito, 1981; Jensen, 1984). Gerlach (1965) described *T. acer* collected from brown algae from a rockpool, in Svalbard. *T. acer* is morphologically and molecularly the closest relative to *T. melnikovi* (Tchesunov, 2006; Timchenko et al., 2024b).

Subsequently, accurate discrimination between closely related species canbe challenging. Morphological variation may arise as a result of different environmental conditions and food availability, a phenomenon known as phenotypic plasticity (West-Eberhard, 1989). Alternatively, cryptic speciation can result in specimens appearing identical but representing genetically distinct populations. The method of fixation can introduce artificial variation, for example through shrinking, as well as differences in how researchers measure morphological characteristics (Mokievsky et al., 2024). Thus, it is imperative that an integrative taxonomic approach, where both morphological and molecular data are evaluated, is taken for taxonomic assignments.

In this study, we used morphological and molecular (amplicon and genomic) data to investigate (i) where *T. melnikovi* persists in Svalbard fjords following landfast ice melt; (ii) whether nematodes occurring on macroalgae can be distinguished morphologically between *T. melnikovi* and *T. acer*; (iii) what the phylogenetic relationship based on 18S rDNA is between specimens from sea-ice and macroalgae; (iv) whether habitat-associated phenotypic variation or cryptic speciation is present in Svalbard populations of *T. melnikovi*.

## Materials and methods

### Study area

Landfast-ice, sediment and macroalgae samples were collected from different fjords around Svalbard: Van Mijenfjorden on the southwest coast of Spitsbergen (Svalbard), Wijdefjorden-Austfjorden in northern Spitsbergen, and Wahlenbergfjorden on Nordaustlandet, north-east Svalbard (Fig. 1). These fjords were chosen, as they had been previously sampled for sea-ice and we wanted to investigate if the summer habitat of sympagic nematodes could be found. Van Mijenfjorden and Wahlenbergfjorden were chosen based on data obtained by Pitusi et al. (2021) & Pitusi et al. (2023). Wijdefjorden-Austfjorden was chosen based on the sea-ice sampling field campaign during 2021 (Duncan et al., 2024b).

**Fig. 1.**
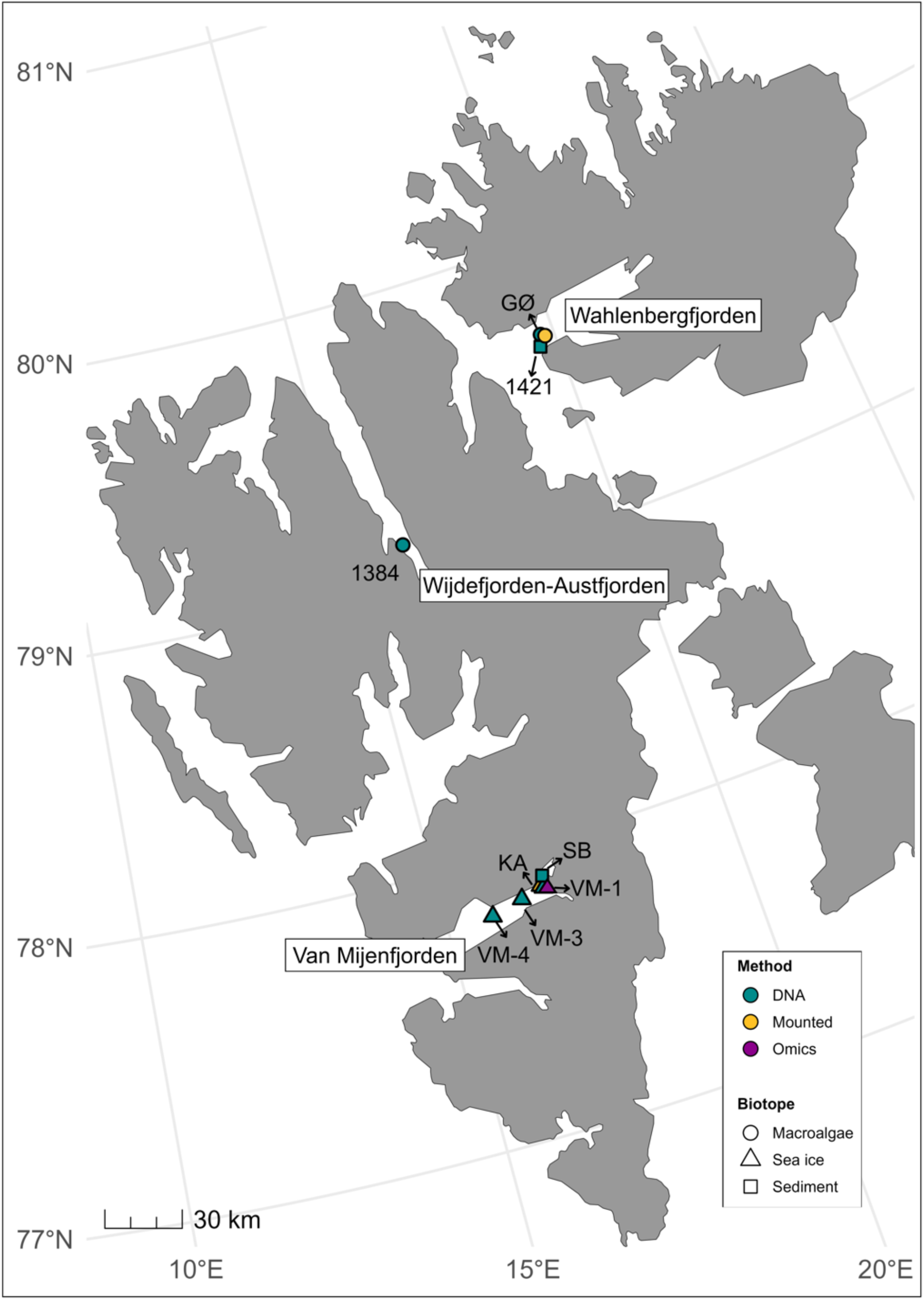
Map of sampling locations for nematodes from macroalgae, sea-ice, and sediment, in Svalbard fjords, with biotope and laboratory methods represented by different shapes and colours. GØ = Gyldenøyane; KA = Kapp Amsterdam; SB = Sveabukta.

Van Mijenfjorden (77.7°N, 15.5°E) is a 70-km-long and 10-km-wide fjord that is partially enclosed at its mouth by an island (Akseløya), regulating the exchange of warm and saline Atlantic water that follows along the shelf via the West Spitsbergen Current (WSC) (Høyland et al. 2009). Consequently, the formation of sea-ice is relatively reliable, as warm Atlantic water typically does not typically enter the fjord until late spring (Duncan et al., 2024c). Wijdefjorden (79.7°N, 15.3°E) is a 110-km long fjord, with the innermost part called Austfjorden (79.1°N, 16.1°E), which is 35-km long and 5-km wide. The presence of a cross-fjord sill (at 60-m) limits water exchange and results in relatively reliable sea-ice cover from January to May (Allaart et al., 2020). The 50-km long and 15-km wide Wahlenbergfjorden (79.7°N, 20.5°E) is situated on Nordaustlandet. This fjord is normally ice-covered in winter but is exposed to Atlantic-derived water at depth from Hinlopen Strait (Bartels et al., 2018).

### Field sampling

Landfast-ice samples (Fig. 2A) were collected in only Van Mijenfjorden in 2015, 2021, 2022, and 2024 (Table 1). Sampling locations were accessed with snowmobiles and sea-ice samples were collected from undisturbed snow with a KOVACS ice corer (ø = 9 cm, Kovacs Ice Coring Equipment, USA). Three ice cores were collected for sympagic nematodes and at each sampling site environmental parameters such as snow depth and ice thickness were recorded. The lower 3 cm of each ice core were collected and they were placed in individual ziplock bags to which 100 mL of 0.7 µm filtered seawater (GF/F, Whatman) was added per 1-cm of sea-ice (Garrison and Buck 1986; Spindler and Dieckmann 1986), before being melted in the dark at +4°C for ∼ 24 h, at the University Centre in Svalbard. For more details on the sampling see Pitusi et al. (2026). The melted sea-ice sections were concentrated over a 20 µm sieve and consequently transferred to a petri dish. A Leica MZ16 stereomicroscope (magnification 0.71-11x) was used to inspect samples, and individual ice nematodes were picked from the sample with a glass Pasteur pipette and transferred to individual Eppendorf tubes. The picked nematodes were preserved in either 4 % formalin or 96 % ethanol to allow for both temporary and permanent mounting, or in *RNAlater* (Thermo Fisher Scientific) for genomic sequencing (Supplementary Data S3).

**Fig. 2.**
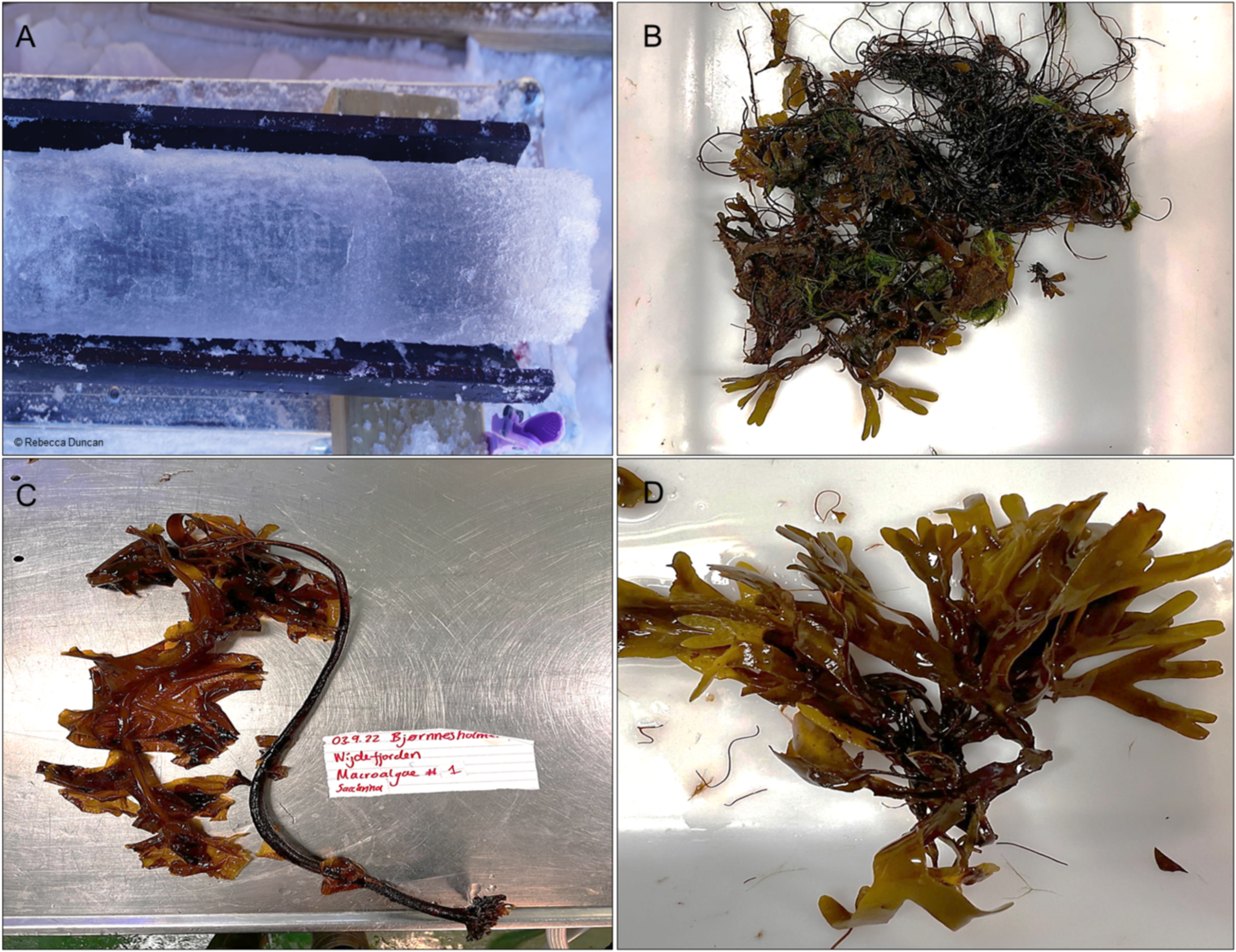
Sampled biotopes for *Theristus*: **A** landfast ice core from Van Mijenfjorden; **B** mixed macroalgae and **C** *Saccharina* sp. retrieved in Wijdefjorden-Austfjorden; **D** *Fucus* sp. sampled at Gyldenøyane.

**Table 1.**
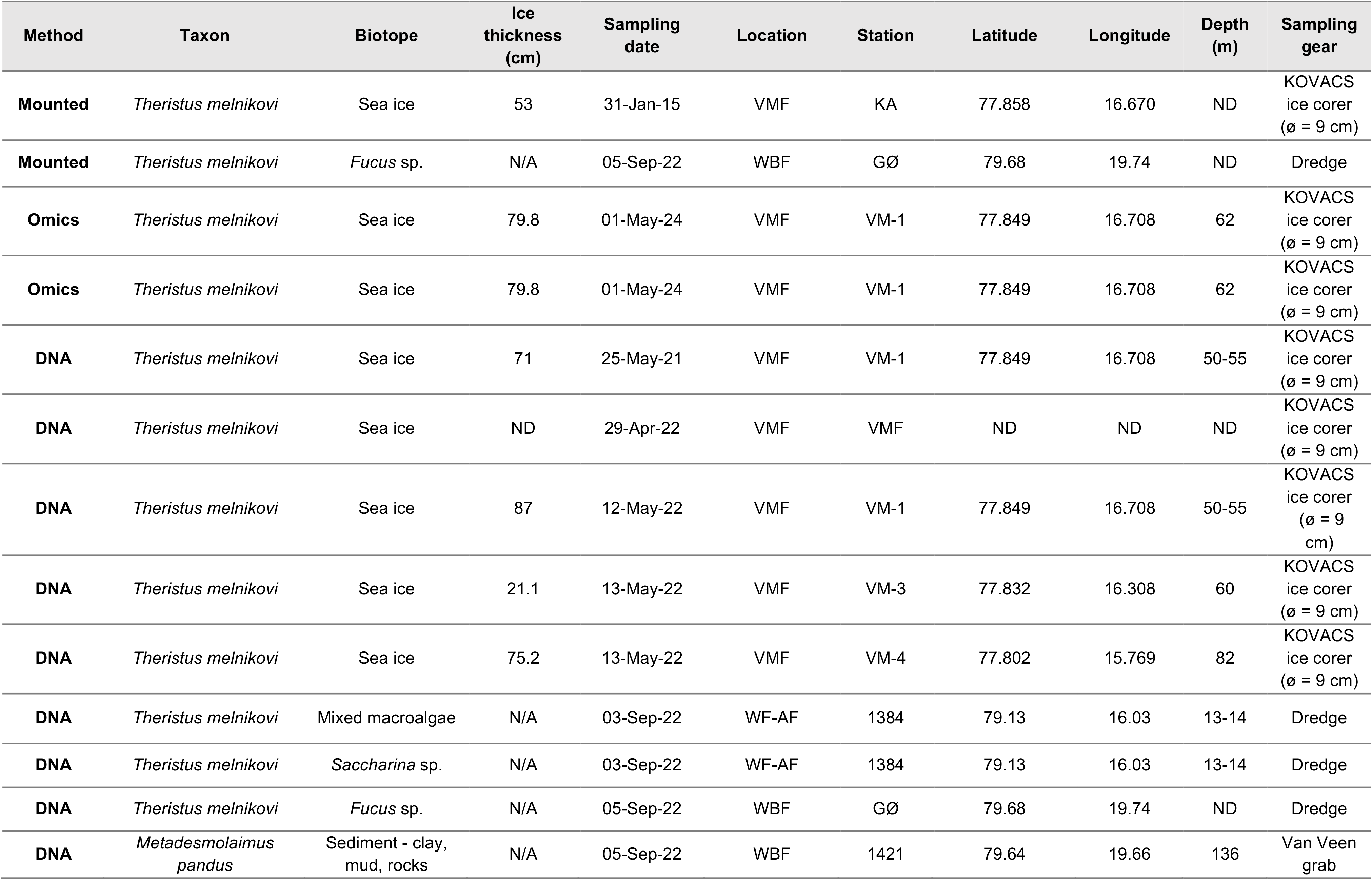
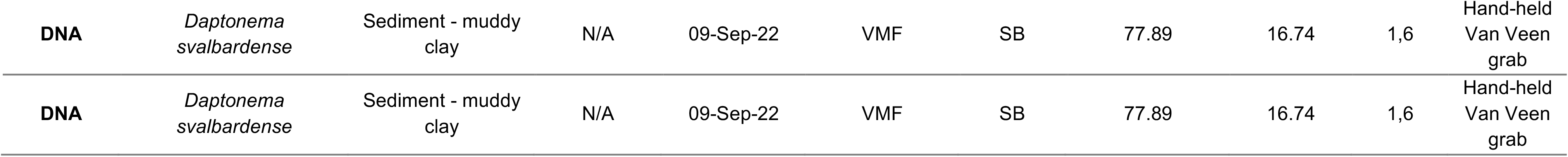
Overview of sampling locations visited in 2015, 2021, 2022, and 2024, from which nematodes were collected. Location abbreviations are as followed – VMF:Van Mijenfjorden, WBF: Wahlenbergfjord, WF-AF: Wijdefjorden-Austfjorden; KA: Kapp Amsterdam; GØ; Gyldénøyane; SB: Sveabukta. N/A = Not applicable; ND = No data.

Sediment and macroalgae samples (Fig. 2B-D) were collected with a Van Veen Grab (KC Denmark, Denmark) and by hand from a Polarcircle, during a cruise onboard of *R/V Helmer Hanssen*, in 2022, from all fjords investigated (Fig. 1). Sediment samples were sieved through a 105 and 44 µm sieve and preserved in 96 % ethanol or 4 % formaldehyde. Macroalgae samples were washed in freshwater, and the water was poured through a 44 µm sieve. The concentrated sample was transferred to a 250 mL sampling bottle and preserved in either 4 % formaldehyde (for taxonomic determination) or 96 % ethanol (molecular analysis).

### Morphological analysis

Formalin-preserved sea-ice samples were concentrated over a 20 μm sieve, from which a total of 15 nematodes of the genus *Theristus* were picked for morphological analysis. The sediment and macroalgae samples were concentrated over a series of sieves 365-160-45 μm, and the concentrates were checked under a stereomicroscope (Olympus SZX12, 3.5-31.5x, Japan). From the macroalgae samples, 24 nematodes of the genus *Theristus* were picked by Dr Oleksandr Holovachov. All nematodes were placed in embryo dishes and processed after an adapted solution from Seinhorst (1966) by placing them in a solution of 4 % glycerol in 50 % ethanol/water for seven days, at room temperature. The glycerol solution was topped up every second day. The permanent slides were prepared by placing nematodes in a drop of glycerol after which the slide was sealed with a paraffin wax ring and clear nail polish. All specimens were identified using Nikon Eclipse 80i (Nikon, Japan) with DIC optics and photographed with a Sony Alpha 6300 (Sony, Japan) mounted on it. Images were processed using ImageJ (Schneider et al., 2012), which was used to obtain various morphometrics such as body length, amphid width, tail length, cephalic setae length and other measurements needed to calculate taxonomic ratios after Hunt & Palomares-Ruis, 2025. Morphological vouchers on permanent glass slides were deposited in the marine invertebrate collection of the Arctic University Museum of Norway (UiT), Tromsø, Norway, under accession numbers TSZY-548-553 (sea-ice) and TSZY-572-575 (macroalgae).

Morphometrics of specimens from Svalbard were compared to records of *Theristus melnikovi* from other Arctic regions, and *Theristus acer* from around the globe (Table 3-6).

### Amplicon sequencing and assembly

In total, 48 nematodes from macroalgae (15), sea-ice (17) and sediment (16), were mounted on temporary microscope slides to be identified and vouchered prior to amplicon sequencing. They were put in drops of MilliQ water and a cover slip was gently placed over the specimen secured with silicone grease (Danco, USA). Depending on the preservation state of the specimen, photographs were taken of the whole body, stoma, amphid, tail, and genitalia (if present) with a Sony Alpha 6300 (Sony, Japan) mounted on either a Nikon Eclipse 80*i* (Nikon, Japan) or Nikon Eclipse N*i* (Nikon, Japan) microscopes equipped with differential interference contrast optics. Photos were taken at 2X, 10X, 20X, 40X, and 60X magnifications. Subsequently, the specimens were placed in new individual Eppendorf tubes with 20-µL QuickExtract DNA Extraction Solution (Lucigen, UK). The samples were extracted at 65°C for 45 min and 98°C for 2 min. Extracts were frozen at -20°C until further processing. After extraction, the whole 18S rRNA fragment was amplified with the following primers. The first fragment was amplified with 988F (5’-CTCAAAGATTAAGCCATGC-3’) or 1096F (5’-GGTAATTCTGGAGCTAATAC-3’) together with 1912R (5’-TTTACGGTCA-GAACTAGGG-3’) (Holterman et al., 2006). The second fragment was amplified with 1813F (5’-CTGCGTGAGAGGTGAAAT-3’) and 2646R (5’-GCTACCTTGGTTACGACTTTT-3’) (Holterman et al., 2006). The primer pairs amplified 962-980 bp and 870-880 bp, respectively. The PCR was performed in a final volume of 25 µL, which consisted of 23 µL mastermix and 2 µL DNA. The mastermix consisted of 2.5 µL 10 × DreamTaq Buffer (Thermo Scientific, USA), 2 μL dNTPS (2.5 mM), 1 μL forward and reverse primers (10 μM), 0.75 μL DreamTaq Polymerase (Thermo Scientific, USA), 1 μL MgCl_2_ (25 nM), and 14.75 μL MilliQ water. The following PCR reaction was used: 94°C – 5 min; 5 x (94°C – 30 sec, 45°C – 30 sec, 72°C -70); 35 x (94°C – 30 sec, 54°C – 30 sec, 72°C – 70 sec); 72°C – 5 min, with an Eppendorf Mastercycler EP Gradient S Thermal Cycler (Eppendorf, Germany). Amplicons were purified using HigherPurity solid-phase reversible immobilization (SPRI) cleaning (Canvax, Córdoba, Spain). During the cleaning process, the amplified DNA of 31 nematodes was lost. Subsequently, amplicons of 17 specimens were sequenced bi-directionally to increase the confidence of the base call along the length of the amplicon. Five of these individuals came from macroalgae, nine from sea-ice, and three from sediment. Sanger sequencing was conducted at Eurofins Genomics (GATC Biotech LightRun, Germany). Sequences were analysed, trimmed, and assembled in Geneious V 2024.0 (Biomatters Ltd New Zealand). The length of successful 18S sequences were 837-1684 bp for *Theristus melnikovi* from sea-ice, 922-1696 bp for *Theristus melnikovi* from macroalgae, and 1690-1692 bp for *Daptonema svalbardense* (Gerlach, 1965) Lorenzen, 1977 from sediment. The sequences were exported and submitted to GenBank under the accession numbers listed in Table 2. All sequences were run through the GenBank database using BLAST with default settings (NCBI, http://www.ncbi.nlm.nih.gov/).

**Table 2.** The DNA sequences obtained in this study with their GenBank accession numbers, isolate number, and sampling location.

| Species | Accession number | Isolate number | Location |
| --- | --- | --- | --- |
| *Theristus melnikovi* | PZ805126 | #22 | Van Mijefjorden, Svalbard |
| *Theristus melnikovi* | PZ805127 | #42 | Van Mijefjorden, Svalbard |
| *Theristus melnikovi* | PZ805128 | #46 | Van Mijefjorden, Svalbard |
| *Theristus melnikovi* | PZ805129 | #52 | Van Mijefjorden, Svalbard |
| *Theristus melnikovi* | PZ805130 | #60 | Van Mijefjorden, Svalbard |
| *Theristus melnikovi* | PZ805131 | #61 | Van Mijefjorden, Svalbard |
| *Theristus melnikovi* | PZ805132 | #63 | Van Mijefjorden, Svalbard |
| *Theristus melnikovi* | PZ805133 | #64 | Van Mijefjorden, Svalbard |
| *Theristus melnikovi* | PZ805134 | #72 | Van Mijefjorden, Svalbard |
| *Theristus melnikovi* | PZ805135 | #79 | Wahlenbergfjorden, Svalbard |
| *Theristus melnikovi* | PZ805136 | #86 | Wahlenbergfjorden, Svalbard |
| *Theristus melnikovi* | PZ805137 | #87 | Wahlenbergfjorden, Svalbard |
| *Theristus melnikovi* | PZ805138 | #90 | Wijdefjorden-Austfjorden, Svalbard |
| *Theristus melnikovi* | PZ805139 | #92 | Wijdefjorden-Austfjorden, Svalbard |
| *Theristus melnikovi* | PZ803337 | GAS | Van Mijefjorden, Svalbard |
| *Theristus melnikovi* | PZ803338 | GAI | Van Mijefjorden, Svalbard |
| *Metadesmolaimus pandus* | PZ805125 | #108 | Wahlenbergfjorden, Svalbard |
| *Daptonema svalbardense* | PZ805123 | #121 | Van Mijefjorden, Svalbard |
| *Daptonema svalbardense* | PZ805124 | #122 | Van Mijefjorden, Svalbard |

In addition, two 18S rRNA sequences were mined from assembled nuclear genomes (Supplementary Data S3) using *barrnap* (Galaxy version 1.2.2) on the usegalaxy.eu server (equivalent to standalone version 0.9, https://github.com/tseemann/barrnap).

### Phylogenetic analysis

All newly generated 18S rRNA sequences (both sequenced directly and mined from genome assemblies) were combined with published sequences of other species from the family Xyalidae, using *Geomonhystera villosa* as an outgroup (see Table S1 for sequences downloaded and used). AliView ver. 1.30 (Larsson, 2014) was used to align sequences and trim them, resulting in a nucleotide matrix that included 70 terminal taxa belonging to 25 species and was 929 bases long (The sequence alignment used for phylogenetic analysis are provided as Supplementary Data S1). The phylogenetic tree was inferred with IQTree ver. 2.3.1-macOS-arm (Minh et al., 2020) using the unpartitioned alignment and built in ModelFinder (Kalyaanamoorthy et al., 2017) with the following command: iqtree2 -s ./input.fas -st DNA -m MFP -b 1000 -T AUTO.

### Haplotype network

The PopART 1.7 software (Clement et al., 2002; Leigh et al., 2015) was used to explore haplotype diversity of newly generated 18S sequences of *T. melnikovi* (Supplementary Data S2) and publicly available in GenBank sequences of *T. melnikovi, T. acer, T. agilis* and *T. ensiferus* (Table S1). AliView ver. 1.30 (Larsson, 2014) was used to align sequences and trim them, resulting in a nucleotide matrix that included 45 sequences and was 860 bases long, providing an input for TCS network.

## Results

### Taxonomic account

Order Monhysterida Filipjev 1929

Family Xyalidae Chitwood 1951

Genus *Theristus* Bastian 1865

*Theristus melnikovi* Tchesunov 1986

### Material examined

Five males and 10 females of sympagic *Theristus melnikovi* (TSZY-548-553, in the collection of the Arctic University Museum of Norway (UiT), Norway) were examined morphologically. Five juveniles, six females and 13 males of macroalgal *T. melnikovi* (TSZY-572-575, in the collection of the Arctic University Museum of Norway (UiT), Norway) were also examined. In addition, seven juveniles, one male and female of sympagic *T. melnikovi* and five juveniles of macroalgal *T. melnikovi* were examined and photographed prior to DNA extraction and consequent destruction.

## Descriptions

### Sympagic Theristus melnikovi

#### Males

Body long and slender, cylindrical, often curved ventrad, more so along the tail in females and in males (Fig. 3A). Cuticle finely striated. Labial region rounded or truncate conoid (Fig. 3B). Inner labial sensilla small setiform, 1.5 μm long, located on the anterior surface of each lip; outer labial and cephalic sensilla setiform, almost similar in length, arranged in a single circle (Fig. 3C); thus, anterior sensilla arranged in a 6+10 pattern. Stoma funnel-shaped overall, with narrow cheilostom, weakly cuticularised broad gymnostom and conoid stegostom gradually extending into pharyngeal lumen, unarmed (Fig. 3B). Amphid located posterior to stoma base, circular with distinct cuticular edge (Fig. 3D), occupying 17–26 % of the corresponding body diameter, with distinct *fusus amphidialis* carrying nerve endings. Renette not observed. Pharynx uniformly cylindrical, evenly muscularised along its entire length. Cardia indistinct. Reproductive system diorchic, anterior testis anteriorly outstretched and located to the left of the intestine, short of cardia; posterior testis reflexed and located to the right of the intestine. Vas deferens small, ejaculatory gland small elongated; ejaculatory duct not visible. Spicules L-shaped, with narrow bifid distal end, curved in the middle and with widened proximal end (Fig. 3E), equal to 0.8–1.2 cloacal body diameters in length. Gubernaculum with broad dorsal apophyses, well-cuticularized, surrounding distal part of spicules. Very small midventral procloacal supplements in the form of thin tubular channels in the cuticle are visible in some individuals, few in number. Tail conical, pointed at the end, strongly curved ventrad. Spinneret present but caudal glands are often indistinct due to fixation.

**Fig. 3.**
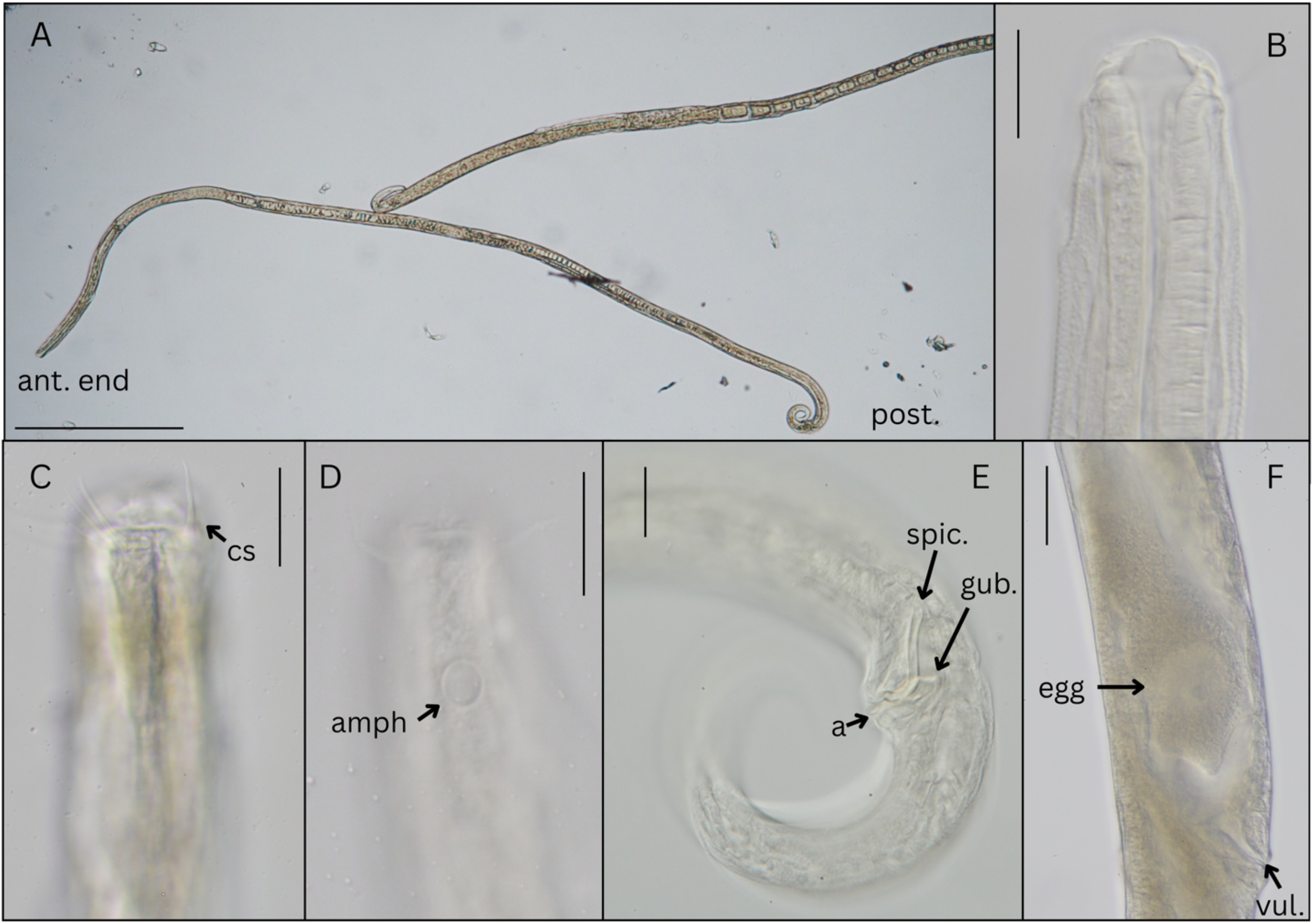
Photomicrographs of *Theristus melnikovi* from Svalbard fjord ice. Details are shown - **A** Specimen on slide with anterior end (ant. end) and posterior end (post. end); **B** Stoma; **C** Anterior end with cephalic setae (cs); **D** Amphid (amph) position and shape; **E** Male cloacal region showing the anal opening (a), spicules (spic.), and gubernaculum (gub.); **F** Vulva (Vul.) and egg of a female. Scale bars: A, C-F = 20 µm; B = 500 µm.

#### Females

Similar to males in most characteristics with some exceptions – larger mid-body diameter with females up to 58 μm compared to 33 μm. Cephalic region more rounded. Reproductive system monodelphic with ovary to the left of the intestine. Uterus is elongated without distinct post vulval uterine sac. Vulvar glands in front and behind the vagina. Intrauterine eggs present with no juveniles inside (Fig. 3F). Vagina oblique, with slightly cuticularized vulva.

#### Juveniles

Morphologically similar to the adult stages, except for the body size and absence of reproductive system. The cephalic region is more rounded rather than conically raised.

### Macroalgal Theristus melnikovi

#### Males

Body elongated and visibly narrowing towards the extremities (Fig. 4A), more fusiform than in the sympagic specimens, curved ventrad along the tail in females and in males. Cuticle finely striated. Labial region rounded with slightly raised lips. Inner labial sensilla small setiform, located on the anterior surface of each lip, outer labial and cephalic sensilla setiform, almost similar in length, arranged in a single circle (Fig. 4B), anterior sensilla arranged in a 6+10 pattern. Stoma relatively small and funnel-shaped (Fig. 4C), with narrow cheilostom, weakly cuticularised broad gymnostom and funnel-shaped stegostom gradually extending into pharyngeal lumen, unarmed. Amphid located posterior to stoma base, circular with distinct cuticular edge (Fig. 4D), occupying 14–21 % of the corresponding body diameter. Pharynx uniformly cylindrical, evenly muscularised along its entire length. Cardia indistinct. Some individuals have somatic setae visible along the body, especially noticeable along the pharyngeal region and on the tail (Fig. 4E), but not clearly observable in all specimens. Reproductive system diorchic, anterior testis anteriorly outstretched and located to the left of the intestine, short of cardia; posterior testis reflexed and located to the right of the intestine. Spicules L-shaped, curved with a kink in the middle, with widened proximal and narrowed distal ends (Fig. 4F), equal to 0.7–1.2 cloacal body diameters in length. Gubernaculum with broad dorsal apophyses, well-cuticularized, surrounding distal part of spicules (Fig. 4G). No precloacal supplements. Tail conical, pointed at the end, strongly curved ventrad. Spinneret present but caudal glands are not visible.

**Fig. 4.**
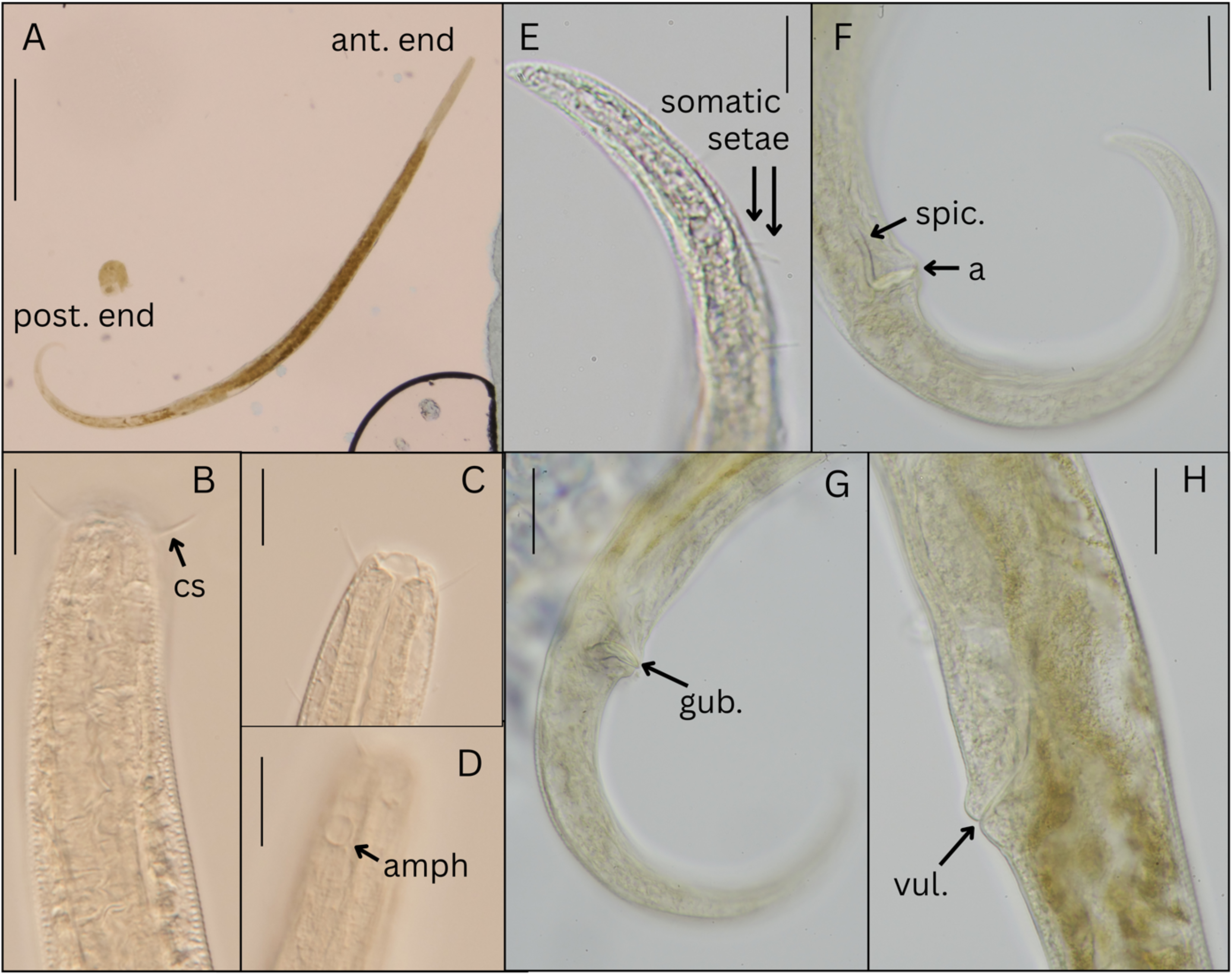
Photomicrographs of *Theristus melnikovi* from Svalbard macroalgae. Details are shown - **A** Specimen on slide with anterior end (ant. end) and posterior end (post. end); **B** Anterior end with cephalic setae (cs); **C** Stoma; **D** Amphid (amph) position and shape; **E** Somatic setae on tail; **F** Male cloacal region showing the anal opening (a) and spicules (spic.); **G** Male cloacal region showing the gubernaculum (gub.); **H** Vulva (Vul.). Scale bars: A = 500 µm; B-H = 20 µm.

#### Females

Similar to males in most characteristics with some exceptions including broader mid-section (52–66 μm vs 30–48 μm). Reproductive system monodelphic with ovary to the left of the intestine. In specimens with a broader mid-section, there appears to be a post vulvar portion while in thinner individuals, there is no post vulvar portion, which could be related to the reproductive age of individual specimens.

#### Juveniles

Morphologically similar to the adult stages, except for the body size and absence of a reproductive system. Somatic setae distinctly visible.

### Morphological comparison

Comparison of both mounted and barcoded specimens of sympagic *Theristus melnikovi* from Svalbard showed that specimens closely matched other Arctic *T. melnikovi* in general morphology and measurements (Table 3 for males; Table 5 for females). Macroalgal specimens resembled *Theristus acer* in overall body form yet retained diagnostic characteristics of *T. melnikovi*.

**Table 3.** Morphometrics (in μm except the ratios a, b, c, c’, and where the percentage (%) is stated) of *Theristus melnikovi* males from different biotopes from this study and other (Arctic) research (Tchesunov (1986); Tchesunov and Portnova (2005); Timchenko et al. (2024b)). The ID corresponds to deposited specimens or GenBank Accession numbers. CAB = Central Arctic Basin; KS = Kara Sea; VMF = Van Mijenfjorden; WBF = Wahlenbergfjorden; WS = White Sea.

| Taxon | *Theristus melnikovi* |  |  |  |  |  |
| --- | --- | --- | --- | --- | --- | --- |
| Location | VMF, Svalbard | VMF, Svalbard | WBF, Svalbard | CAB | WS | KS |
| ID | TSZY-548; TSZY-552-553 | PZ805130 | TSZY-572-575 | SP-22 TV-1981 ice 2-1 | – | 6234//1 |
| Biotope | Landfast ice | Landfast ice | *Fucus* sp. | Perennial ice | Coastal sea ice | First-year ice |
| Reference | This study | This study | This study | Tchesunov (1986) | Tchesunov and Portnova (2005) | Timchenko et al. (2024b) |
| Sampling date | Jan-15 | Apr-22 | Sep-22 | Apr-81 | Mar-97 | Jul-19 |
| *n* | 5 | 1 | 13 | 4 | 1 | 1 |
| Total body length | 2584 ± 212 (2356-2857) | 3088.6 | 1855 ± 172 (1523-2209) | 3150 (2060–3440) | 2500 | 2458 |
| Body diameter at midbody | 33 ± 1.2 (32-35) | 35.5 | 39 ± 5 (30-48) | 50 (28–47) | 32 | 29 |
| Pharynx length | 425 ± 332 (378-452) | 449.0 | 287 ± 58 (152-408) | 502.8 | 367.6 | 305 |
| Tail length | 199 ± 272 (167-249) | 217.4 | 215 ± 19 (177-241) | 222 (171–233) | 247.5 | 187 |
| Body diameter at anal | 31 ± 2.2 (28-35) | 41.5 | 31 ± 3 (24-37) | 41 (30–37) | 33 | 26 |
| a | 78.2 ± 5.6 (72.2-87.7) | 87.1 | 47.9 ± 4.9 (41-61) | 63.4 (68.6–79.8) | 78.1 | 84 |
| b | 6.1 ± 0.3 (5.6-6.3) | 6.9 | 6.8 ± 1.6 (4.5-11.3) | 6.3 (5.68–7.52) | 6.8 | 8.1 |
| c | 13.8 ± 1.8 (12.4-17.1) | 14.2 | 8.6 ± 0.6 (7.6-9.7) | 14.2 (12.4–16.1) | 10 | 13.2 |
| c' | 6.2 ± 0.7 (4.8-6.7) | 5.2 | 7.1 ± 1.0 (5.7-10) | 5.4 (5.7–6.2) | 7.5 | 7.2 |
| **Labial region diameter** | 14 ± 2.1 (10.6-15.8) | 13.0 | 11.4 ± 1.6 (8.1-13.6) | ? | ? | ? |
| **Cephalic setae length** | 11.8 ± 1.4 (9.3-13.2) | 16.1 | 7.7 ± 1.6 (5.1-10.7) | 15 (15–16) | 12 | 11 |
| **CSL/LRD** | 0.9 ± 0.2 (0.8-1.2) | 1.2 | 0.7 ± 0.1 (0.5-0.9) | ? | ? | ? |
| **Amphidial fovea width** | 5.6 ± 1 (4.1-6.6) | 5 | 4.3 ± 0.3 (3.6-4.9) | 8 | 8.5 | 6 |
| **Amphidial fovea from anterior end** | 29 ± 1.7 (26.2-30.6) | 32.3 | 18.3 ± 2 (16.0-22.5) | 36 (31–37) | 43 | 28 |
| **Head diameter** | 22 ± 1.5 (19-23) | 22.9 | 19 ± 1.5 (15-21) | 22 (21–23) | 21 | 19 |
| **cbd** | 25 ± 0.9 (24-26) | 26.5 | 23 ± 2 (21-28) | 30 | ? | 22 |
| **Amphidial fovea width/cbd (%)** | 19 ± 2.8 (15-22) | 19 | 19 (17-18) | 26 (35–38) | 35 | 28 |
| **Stoma length** | 6.5 ± 0.9 (5.5-7.8) | 7.6 | 4.3 ± 0.7 (3.1-5.4) | 14 | 13 | 8 |
| **Stoma width** | 9.9 ± 1.1 (8.5-11) | 8.8 | 7.6 ± 0.7 (6.4-8.5) | 13 | 12.5 | 12 |
| **Rectum length** | 20 ± 1.2 (18-23) | 16.9 | 20 ± 1 (17-22) | ? | ? | ? |
| **Spicule length along arc** | 33 ± 3.2 (28-37) | 34.3 | 30 ± 4.6 (22-37) | 31 | 31 | 36 |
| **Spicule length along chord** | 27 ± 2.2 (23-30) | 29.8 | 24 ± 3.5 (17-29) | 25 (28–31) | 24 | 32 |
| **Gubernaculum apophysis length** | 18 ± 1.2 (15-20) | ? | 18 ± 0.9 (16-19) | 12 (10–13) | 6 | 11 |
| **Somatic setae** | - | - | 5.5 | 6 | ? | 3 |
| **Cloaca to supplement** | ? | ? | ? | 35 | – | – |
| **Supplement to supplement** | ? | ? | ? | 48 | – | – |

$n$ = number of specimens, $a$ = total body length/maximum body diameter, $b$ = total body length/pharynx length, $c$ = total body length/tail length, $c'$ = tail length/body diameter at anal, $cbd$ = corresponding body diameter, $V$ (%) = (vulva from anterior end/total body length) x 100, $V'$ (%) = (vulva from anterior end/anus from anterior end) x 100, - = absence of characteristic, ? = not possible to measure due to body deformation or incorrect positioning.

**Table 4.** Morphometrics (in μm except the ratios a, b, c, c’, and where the percentage (%) is stated) of *Theristus acer* males from intertidal habitats around the globe (Bastian, 1865; Wieser, 1959; Gerlach, 1965; Hopper, 1969; Kito, 1981; Warwick et al., 1998).

| **Taxon** | ***Theristus acer*** |  |  |
| --- | --- | --- | --- |
| **Location** | **Atlantic** | **Pacific** | **Arctic** |
| **Biotope** | **Marine mud** | **Beach sediment & macroalgae** | ***Fucus sp.* - rockpool** |
| **Reference** | **Bastian (1865), Hopper (1969) and Warwick et al. (1998)** | **Wieser (1959) & Kito (1981)** | **Gerlach (1965)** |
| ***n*** | 3 | 6 | 2 |
| **Total body length** | 1600-2390 | 1195-2220 | 1650-1655 |
| **Body diameter at midbody** | 37-58.5 | 26-33 | 42-50 |
| **Pharynx length** | 211-347 | ND | 270-300 |
| **Tail length** | 178-252 | 141-288 | 210-215 |
| **Body diameter at anal** | 38-43 | 34-36 | 35-40 |
| **a** | 28-56.9 | 44.5 | 33-39 |
| **b** | 5-8 | 8.6 | 5.5-6.1 |
| **c** | 8.9-10.4 | 12.7 | 7.7-7.9 |
| **Cephalic setae length** | 10-15 | 8-12 | 13-14 |
| **Amphidial fovea width** | 6-8 | 7-7.5 | 8-9 |
| **Amphididial fovea from anterior end** | ND | ND | 19-28 |
| **Head diameter** | 17-21 | 13-25 | 23 |
| **cbd** | ND | ND | 25-28.1 |
| **Amphidial fovea width/cbd (%)** | ND | 30-42 | ND |
| **Spicule length** | 35-54 | 25-28 | 26 |
| **Gubernaculum apophysis length** | ND | 15-23 | ND |

*n* = number of specimens, a = total body length/maximum body diameter, b = total body length/pharynx length, c = total body length/tail length, cbd = corresponding body diameter, V (%) = (vulva from anterior end/total body length) x 100, ND = no data (Table 4).

**Table 5.**
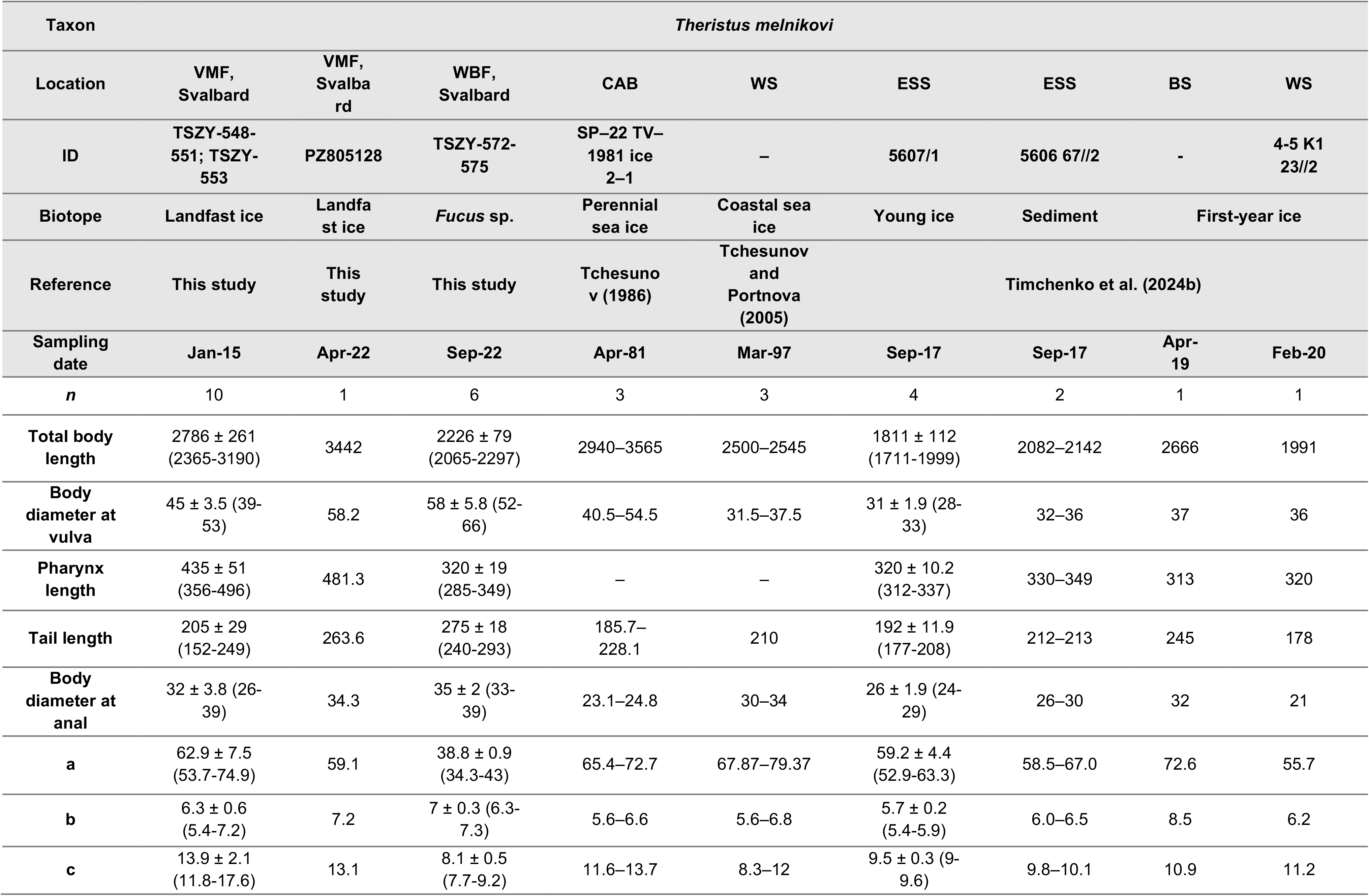
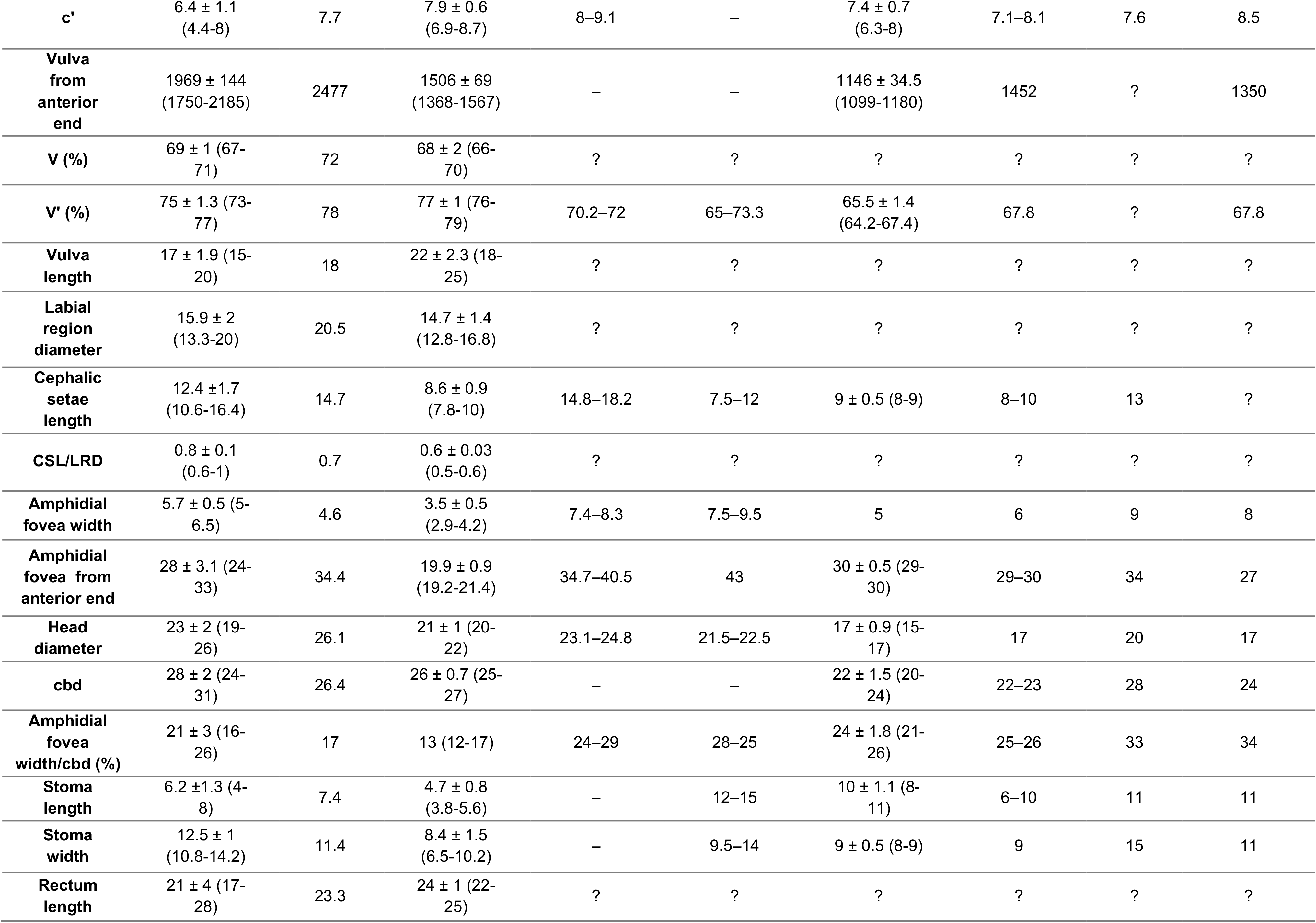
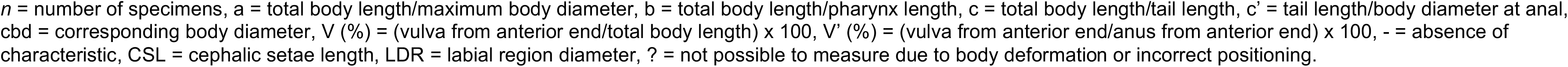
Morphometrics (in μm except the ratios a, b, c, c’, and where the percentage (%) is stated) of *Theristus melnikovi* females from different biotopes from this study and other Arctic research (Tchesunov (1986); Tchesunov and Portnova (2005); Timchenko et al. (2024b)). The ID corresponds to deposited museum specimens or GenBank Accession numbers. BS = Barents Sea; CAB = Central Arctic Basin; ESS = East Siberian Sea; VMF = Van Mijenfjorden; WBF = Wahlenbergfjorden; WS = White Sea.

| Taxon | *Theristus melnikovi* |  |  |  |  |  |  |  |  |
| --- | --- | --- | --- | --- | --- | --- | --- | --- | --- |
| Location | VMF, Svalbard | VMF, Svalbard | WBF, Svalbard | CAB | WS | ESS | ESS | BS | WS |
| ID | TSZY-548-551; TSZY-553 | PZ805128 | TSZY-572-575 | SP-22 TV-1981 ice 2-1 | – | 5607/1 | 5606 67//2 | - | 4-5 K1 23//2 |
| Biotope | Landfast ice | Landfast ice | *Fucus* sp. | Perennial sea ice | Coastal sea ice | Young ice | Sediment | First-year ice |  |
| Reference | This study | This study | This study | Tchesunov (1986) | Tchesunov and Portnova (2005) | Timchenko et al. (2024b) |  |  |  |
| Sampling date | Jan-15 | Apr-22 | Sep-22 | Apr-81 | Mar-97 | Sep-17 | Sep-17 | Apr-19 | Feb-20 |
| *n* | 10 | 1 | 6 | 3 | 3 | 4 | 2 | 1 | 1 |
| Total body length | 2786 ± 261 (2365-3190) | 3442 | 2226 ± 79 (2065-2297) | 2940–3565 | 2500–2545 | 1811 ± 112 (1711-1999) | 2082–2142 | 2666 | 1991 |
| Body diameter at vulva | 45 ± 3.5 (39-53) | 58.2 | 58 ± 5.8 (52-66) | 40.5–54.5 | 31.5–37.5 | 31 ± 1.9 (28-33) | 32–36 | 37 | 36 |
| Pharynx length | 435 ± 51 (356-496) | 481.3 | 320 ± 19 (285-349) | – | – | 320 ± 10.2 (312-337) | 330–349 | 313 | 320 |
| Tail length | 205 ± 29 (152-249) | 263.6 | 275 ± 18 (240-293) | 185.7–228.1 | 210 | 192 ± 11.9 (177-208) | 212–213 | 245 | 178 |
| Body diameter at anal | 32 ± 3.8 (26-39) | 34.3 | 35 ± 2 (33-39) | 23.1–24.8 | 30–34 | 26 ± 1.9 (24-29) | 26–30 | 32 | 21 |
| a | 62.9 ± 7.5 (53.7-74.9) | 59.1 | 38.8 ± 0.9 (34.3-43) | 65.4–72.7 | 67.87–79.37 | 59.2 ± 4.4 (52.9-63.3) | 58.5–67.0 | 72.6 | 55.7 |
| b | 6.3 ± 0.6 (5.4-7.2) | 7.2 | 7 ± 0.3 (6.3-7.3) | 5.6–6.6 | 5.6–6.8 | 5.7 ± 0.2 (5.4-5.9) | 6.0–6.5 | 8.5 | 6.2 |
| c | 13.9 ± 2.1 (11.8-17.6) | 13.1 | 8.1 ± 0.5 (7.7-9.2) | 11.6–13.7 | 8.3–12 | 9.5 ± 0.3 (9-9.6) | 9.8–10.1 | 10.9 | 11.2 |
| **c'** | 6.4 ± 1.1<br>(4.4-8) | 7.7 | 7.9 ± 0.6<br>(6.9-8.7) | 8–9.1 | – | 7.4 ± 0.7<br>(6.3-8) | 7.1–8.1 | 7.6 | 8.5 |
| **Vulva from anterior end** | 1969 ± 144<br>(1750-2185) | 2477 | 1506 ± 69<br>(1368-1567) | – | – | 1146 ± 34.5<br>(1099-1180) | 1452 | ? | 1350 |
| **V (%)** | 69 ± 1 (67-71) | 72 | 68 ± 2 (66-70) | ? | ? | ? | ? | ? | ? |
| **V' (%)** | 75 ± 1.3 (73-77) | 78 | 77 ± 1 (76-79) | 70.2–72 | 65–73.3 | 65.5 ± 1.4<br>(64.2-67.4) | 67.8 | ? | 67.8 |
| **Vulva length** | 17 ± 1.9 (15-20) | 18 | 22 ± 2.3 (18-25) | ? | ? | ? | ? | ? | ? |
| **Labial region diameter** | 15.9 ± 2<br>(13.3-20) | 20.5 | 14.7 ± 1.4<br>(12.8-16.8) | ? | ? | ? | ? | ? | ? |
| **Cephalic setae length** | 12.4 ± 1.7<br>(10.6-16.4) | 14.7 | 8.6 ± 0.9<br>(7.8-10) | 14.8–18.2 | 7.5–12 | 9 ± 0.5 (8-9) | 8–10 | 13 | ? |
| **CSL/LRD** | 0.8 ± 0.1<br>(0.6-1) | 0.7 | 0.6 ± 0.03<br>(0.5-0.6) | ? | ? | ? | ? | ? | ? |
| **Amphidial fovea width** | 5.7 ± 0.5 (5-6.5) | 4.6 | 3.5 ± 0.5<br>(2.9-4.2) | 7.4–8.3 | 7.5–9.5 | 5 | 6 | 9 | 8 |
| **Amphidial fovea from anterior end** | 28 ± 3.1 (24-33) | 34.4 | 19.9 ± 0.9<br>(19.2-21.4) | 34.7–40.5 | 43 | 30 ± 0.5 (29-30) | 29–30 | 34 | 27 |
| **Head diameter** | 23 ± 2 (19-26) | 26.1 | 21 ± 1 (20-22) | 23.1–24.8 | 21.5–22.5 | 17 ± 0.9 (15-17) | 17 | 20 | 17 |
| **cbd** | 28 ± 2 (24-31) | 26.4 | 26 ± 0.7 (25-27) | – | – | 22 ± 1.5 (20-24) | 22–23 | 28 | 24 |
| **Amphidial fovea width/cbd (%)** | 21 ± 3 (16-26) | 17 | 13 (12-17) | 24–29 | 28–25 | 24 ± 1.8 (21-26) | 25–26 | 33 | 34 |
| **Stoma length** | 6.2 ± 1.3 (4-8) | 7.4 | 4.7 ± 0.8<br>(3.8-5.6) | – | 12–15 | 10 ± 1.1 (8-11) | 6–10 | 11 | 11 |
| **Stoma width** | 12.5 ± 1<br>(10.8-14.2) | 11.4 | 8.4 ± 1.5<br>(6.5-10.2) | – | 9.5–14 | 9 ± 0.5 (8-9) | 9 | 15 | 11 |
| **Rectum length** | 21 ± 4 (17-28) | 23.3 | 24 ± 1 (22-25) | ? | ? | ? | ? | ? | ? |

$n$ = number of specimens, $a$ = total body length/maximum body diameter, $b$ = total body length/pharynx length, $c$ = total body length/tail length, $c'$ = tail length/body diameter at anal, $cbd$ = corresponding body diameter, $V$ (%) = (vulva from anterior end/total body length) × 100, $V'$ (%) = (vulva from anterior end/anus from anterior end) × 100, - = absence of characteristic, CSL = cephalic setae length, LDR = labial region diameter, ? = not possible to measure due to body deformation or incorrect positioning.

**Table 6.**
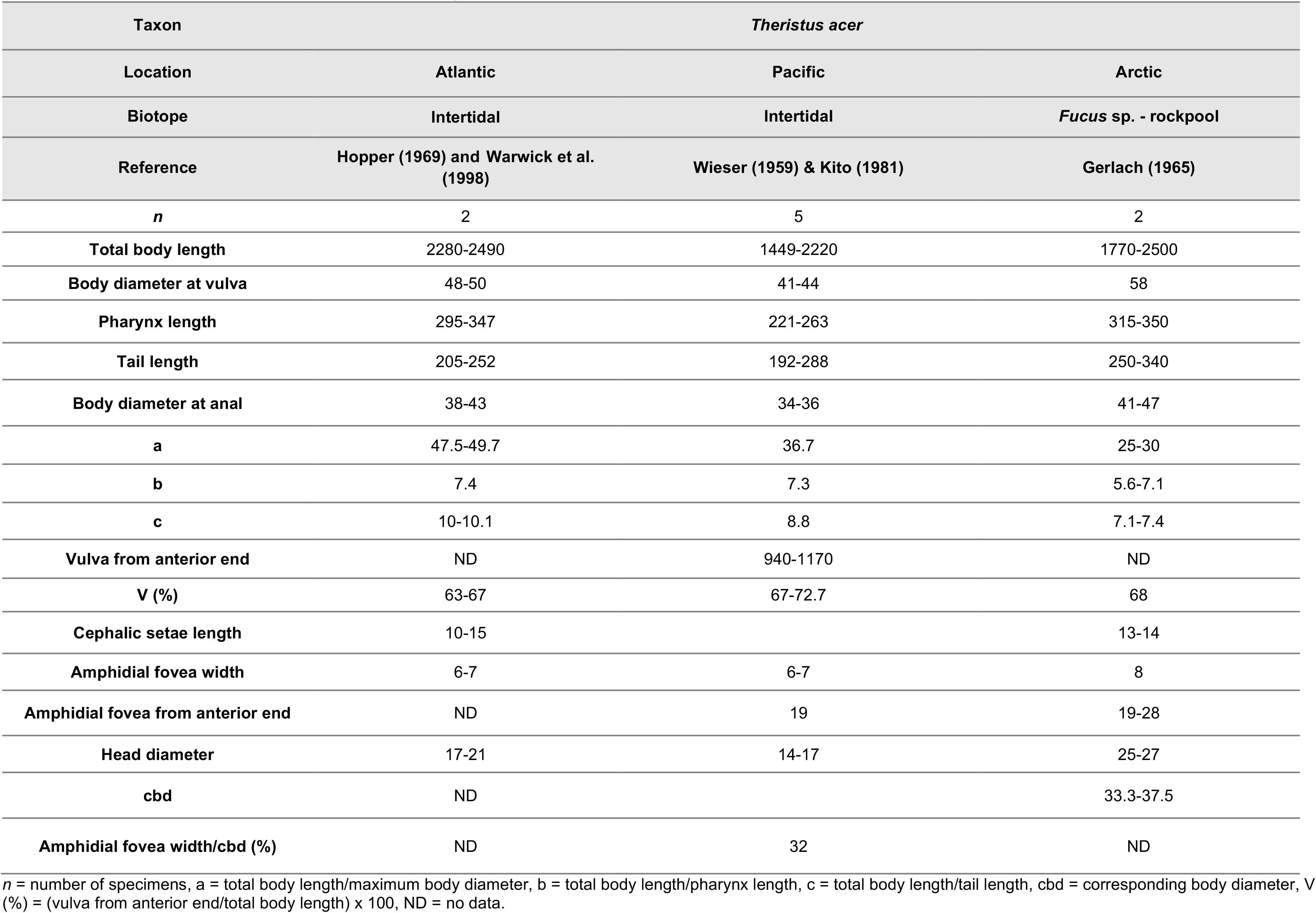
Morphometrics (in μm except the ratios a, b, c, c’, and where the percentage (%) is stated) of *Theristus acer* females from intertidal habitats around the globe (Wieser, 1959; Gerlach, 1965; Hopper, 1969; Kito, 1981; Warwick et al., 1998).

*Theristus acer* is the most morphologically and genetically similar species to *T. melnikovi*, both based on current observations and published data (Tchesunov, 2006; Timchenko et al., 2024b). At the same time, it is a common and broadly distributed species in the North Atlantic (both east and west), with additional records from the North Pacific (Wieser, 1959; Kito, 1981; unpublished 18S rRNA), and one record from brown algae from Svalbard (Gerlach, 1965). Unfortunately, not all species records are backed by detailed morphological and/or molecular data, and the taxonomic status of some populations is questionable. The original description of *T. acer*, based on specimens from the Falmouth estuary (Cornwall, UK), is very brief (Bastian, 1865) (Table 4 for males; Table 5 for females) and, not unexpectedly, does not meet current taxonomic standards, and the type material is inaccessible. As such, we consider the subsequent re-description by Warwick et al. (1998) and sequence data published by Bhadury et al. (2006) from several localities near Plymouth (Devon, UK) to be the most reliable representation of this species and used them to establish the identity of this species. At the same time, specimens described by Gerlach (1965) as *T. acer* from brown algae from Svalbard can possibly represent the non-sympagic *T. melnikovi.* Therefore, the comparison below treats *T. acer* sensu Gerlach, 1965 separately from the North Atlantic and North Pacific populations of the species.

Males of the sympagic *Theristus melnikovi* population from Svalbard were generally consistent with published descriptions from other Arctic localities, including the Central Arctic Basin (CAB), White Sea (WS), and the Kara Sea (KS) (Table 3). On average, Svalbard males were shorter than the individuals recorded from the CAB but overlapped in body length with males from the WS and KS. The exception was the singular individual collected in April 2022, which was similar in body length to the CAB specimens. Svalbard males differed from other Arctic populations in their distinctly smaller stoma, with a mean stomal width and length of 9.9 and 6.5 μm, respectively, compared to 8-14 μm and 12-13 μm. The gubernaculum of Svalbard specimens was longer, averaging 18 μm versus 12 μm in the CAB, 6 μm in the WS, and 11 μm in the KS.

Males from the brown macroalgae *Fucus* in Wahlenbergfjorden (WBF) differed morphometrically from the sympagic *T. melnikovi* populations, particularly in their shorter and distinctly stouter body and smaller anterior structures. WBF males were 1.4 times shorter than individuals from sea-ice in Van Mijenfjorden (VMF) and other Arctic ice-associated populations. WBF males had a relatively broad body diameter (30–48 μm), overlapping with the CAB population (28–47 μm). Body diameter in WBF was 1.4-1.7 times greater than in VMF (32–35 μm), WS (32 μm), and KS (29 μm). In several general morphometric characters, WBF males overlapped with published records of *Theristus acer* from the Arctic, Pacific, and Atlantic. (Table 4). Males of *T. acer* from Svalbard brown algae described by Gerlach (1965) were even shorter and stouter (a = 33–39) than WBF specimens (a = 41–61) but overlapped in their proportional tail length. Both the Atlantic and Pacific *T. acer* populations overlapped with WBF in general measurements such as body length and tail length (Table 3-4). However, other values such as amphid width were larger for the Atlantic (6-7 μm) and Pacific (7–7.5 μm) compared to WBF (3.6–4.9 μm). On average, the Atlantic specimens were also broader (47.9 μm) and more slender (a = 42.5) than the WBF males (39 μm; a = 47.9). Male genitalia was less informative. Spicule and gubernaculum measurements overlapped broadly between WBF and sympagic *T. melnikovi* males despite their differences in body proportions. Comparisons with published *T. acer* populations were limited, as descriptions did not explicitly specify whether the arc or chord length was measured. Nevertheless, the reported values showed substantial variation among geographic populations. For Arctic and Pacific specimens, midventral procloacal supplements were observed, which were also present in the ice-associated *T. melnikovi* from Svalbard, but not the WBF males.

As in the males, the female VMF population falls within the published description of *T. melnikovi* from other Arctic localities (Table 5). Some variation in morphometric measurements was observed, but these differences remained within the range reported for the species.

Females from the brown macroalgae *Fucus* in WBF were shorter (2065–2297 μm) and stouter than sympagic *T. melnikovi* from VMF, the WS, and the Barents Sea (BS). WBF females overlapped in mean body diameter with the singular female from VMF from April 2022 (58.4 μm), the CAB population (40.5–54.5 μm). Mean body length in WBF females (2226 μm) was greater than values reported from sea-ice in the East Siberian Sea (ESS;1811 μm) and the WS (1991 μm) but was comparable with *T. melnikovi*s in ESS sediment (2082–2142 μm). WBF females also had relatively longer tails, reflected in their lower c values (7.7–9.2) compared to sympagic *T. melnikovi* females from VMF (11.8–17.6) and CAB (11.6–13.7). Additionally, WBF females had the smallest stoma, with a length of 3.8-5.6 μm and width of 6.5–10.2 μm. These values most closely overlapped those reported for *T. melnikovi* in ESS sediment (length 6–10 μm; width 9 μm) butwere smaller than those documented from the WS (length 11–15 μm; width 9.5–14 μm). Sympagic *T. melnikovi* and WBF females vulval position overlapped substantially with other ice populations. The singular individual of *T. melnikovi* had the most posterior vulval position (V’ = 78 %) followed by WBF with 77 % and VMF January 2015 with 74 %. Other populations had comparable (CAB = 70.2–72 %) or lower values (e.g. 65.5 in ESS sea-ice).

WBF females overlapped with published records of *Theristus acer* from Arctic, Pacific, and Atlantic localities in various general morphometric characters (Table 5-6). All populations of *T. acer* overlapped with WBF females in regard to their low c values. The Svalbard population described by Gerlach (1965) overlapped with WBF females in body diameter (58 μm vs 52–66 μm for WBF females), pharynx (333 μm vs 320 μm) and tail (295 μm vs 275 μm) length. However, WBF females were considerably more slender (a = 34.3–43 vs 25–30) than *T. acer* from Svalbard. Pacific and Atlantic *T. acer* populations also showed overlap in general measurements such as pharynx and tail length with WBF females. However, Atlantic specimens were more slender (a = 47.5–49.7), whereas WBF females had smaller amphids (2.9–4.2 μm) compared to the *T. acer* populations (6-8 μm).

Juveniles from macroalgae collected in WBF and Wijdefjorden-Austfjorden (WF-AF) were minimally longer than those from sea-ice (1219–2144 μm vs 1138–2064 μm), although body length overlapped considerably. The range in body length most likely can be attributed to different juvenile stages. As observed with the adults, macroalgal juveniles were stouter than the sea-ice individuals (29–49 μm vs 30–38 μm). They also had relatively longer tails, reflected in the lower c values (6.6–9.6 μm vs 9–11.4 μm). Other morphometric measurements, such as head diameter, showed considerable overlap between the two populations (Table S2).

Overall, all life stages from WBF represented a distinctly stout-bodied, long-tailed morphotype that can be regarded as an intermediate morphology between sympagic *T. melnikovi* and *T. acer*.

### Molecular analysis

The phylogenetic analysis and haplotype network revealed limited 18S rRNA sequence variation among sampled Arctic *Theristus melnikovi* populations, while clearly separating them from the included *T. acer* sequences. Despite their morphological resemblance to *T. acer*, all sequenced macroalgal specimens grouped with *T. melnikovi* and clustered most closely with a singular *T. melnikovi* from the White Sea (19TMWS; ON153225).

#### Phylogenetic analysis

Phylogenetic reconstruction using Maximum Likelihood (ML) revealed that all included in the analysis species of the genus *Theristus* separated into five clades (Fig. 5). Clade III is well-supported and monophyletic, composed exclusively of *Theristus melnikovi*, and includes both newly generated (Table 2) and previously published 18S sequences from Svalbard (Pitusi et al., 2021), the Kara and White Sea (Timchenko et al., 2024b). Within this clade, most of the included samples create a broad polytomy with low or no divergence, and nine of the newly obtained sequences (PZ805126-34; PZ80513337-38) are part of this polytomy alongside the sequences from the Kara and White Sea. The five sequences from *T. melnikovi* from macroalgae (PZ805135-38), on the other hand, form a distinct strongly supported clade together with one sequence (ON153214) from the White Sea rather than being situated within or near Clade II of *Theristus acer*. Instead, *T. acer* from the (UK, Netherlands and China) and *T. melnikovi* formed strongly supported but well-defined sister clades, which is also supported by the phylogeny based on *COX1* sequences (Supplementary Data S3; Fig. S3-4). This sister relationship was consistent with the morphological similarity between the species but demonstrated that the macroalgal specimens did not belong to *T. acer*. Notably, several of the newly generated *T. melnikovi* sequences (e.g. PZ805126) had *T. acer* sequences (e.g. AJ966505 and OQ396745) among their closest GenBank matches using BLAST, but phylogenetic analysis using ML securely placed them within the *T. melnikovi* clade.

**Fig. 5.**
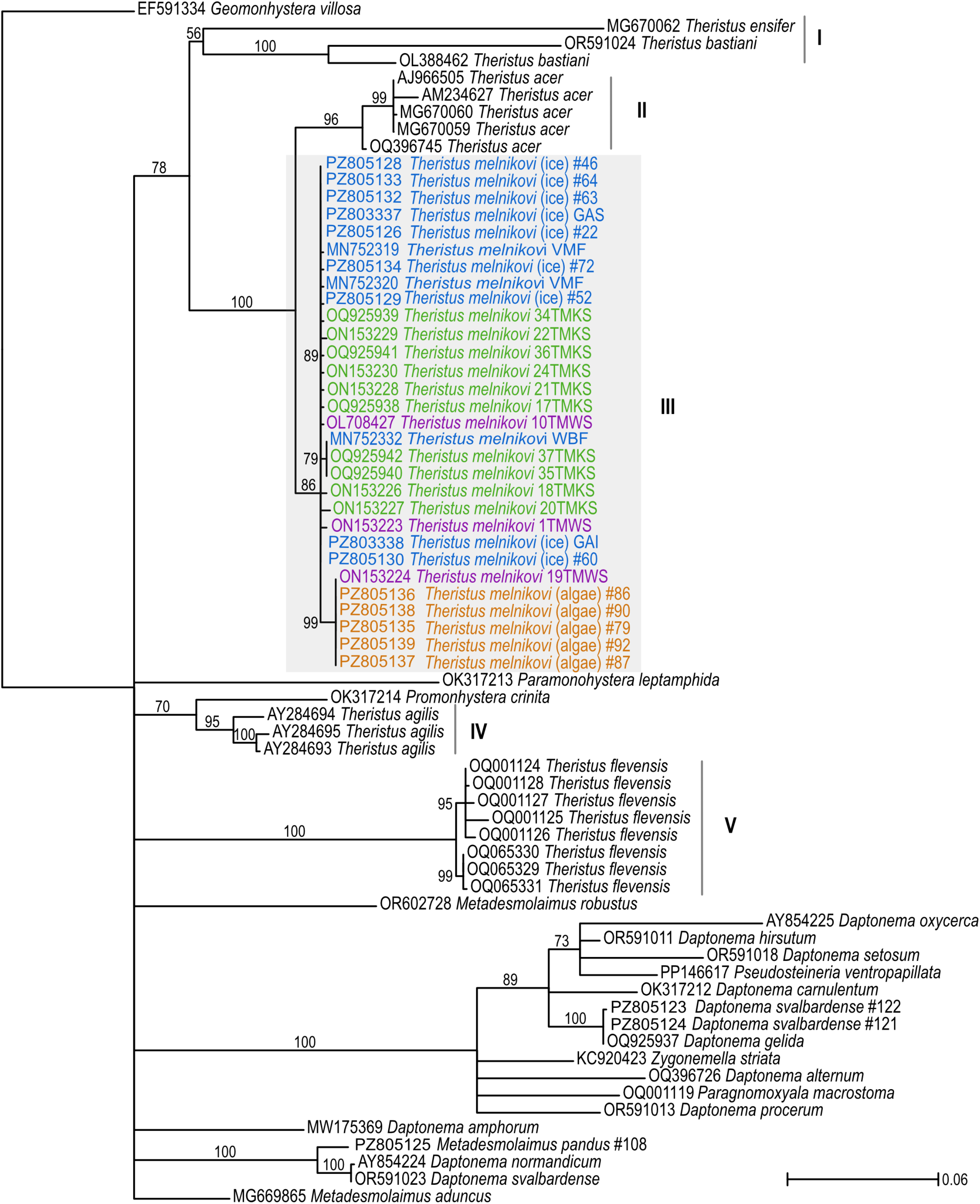
Maximum Likelihood tree inferred from 18S rDNA sequences based on species of the Family Xyalidae with representatives of *Theristus melnikovi* from Svalbard and other Arctic regions. *Geomonhystera villosa* (EF591334) is used as the outgroup. The branches support values: ≥ 0.75 good to moderate support, and ≥ 0.90 strong support. Five *Theristus* clades are labelled with sequences from *Theristus melnikovi* highlighted in a grey box. Sequences used are colour coded: purple for the White Sea, green for the Kara Sea, blue for Svalbard sea-ice, and orange for the Svalbard macroalgae.

#### Haplotype network

The TCS haplotype network based on partial 18S rDNA networks from *Theristus melnikovi* (both sympagic and macroalgal) and *Theristus acer* with the inclusion of two distantly related species as outgroups, *Theristus flevensis* Schuurmans Stekhoven, 1935 and *Theristus agilis* (de Man, 1880) Filipjev, 1918 revealed 11 haplotypes (Figure 6). The Arctic sequences of *T. melnikovi* formed the first three haplotypes where Haplotype 1 (H1) formed the largest node with a total of 23 individuals from the Kara Sea (TMKS), the White Sea (TMWS), and Svalbard (specifically Wahlenbergfjorden and Van Mijenfjorden). Haplotype 2 (H2) was represented by a singular individual of *T. melnikovi* (1TMWS) from the White Sea and was separated from H1 by two mutational steps. Haplotype 3 (H3) was separated from H1 by four mutational steps and included six individuals consisting of five *T. melnikovi* from macroalgae in Svalbard and one *T. melnikovi* from sea-ice from the White Sea (19TMWS). The Arctic haplotypes were separated from the European (MG670000, AJ966505, AM234627) and Chinese (OQ396745) *T. acer* by more than 21 mutational steps. Thus, the macroalgal phenotype that resembled *T. acer* did not have an 18S haplotype affinity to *T. acer.* The distantly related *T. flevensis* and *T. agilis* formed the last haplotypes and were separated from the other *Theristus* species by more than 49 mutation steps.

**Fig. 6.**
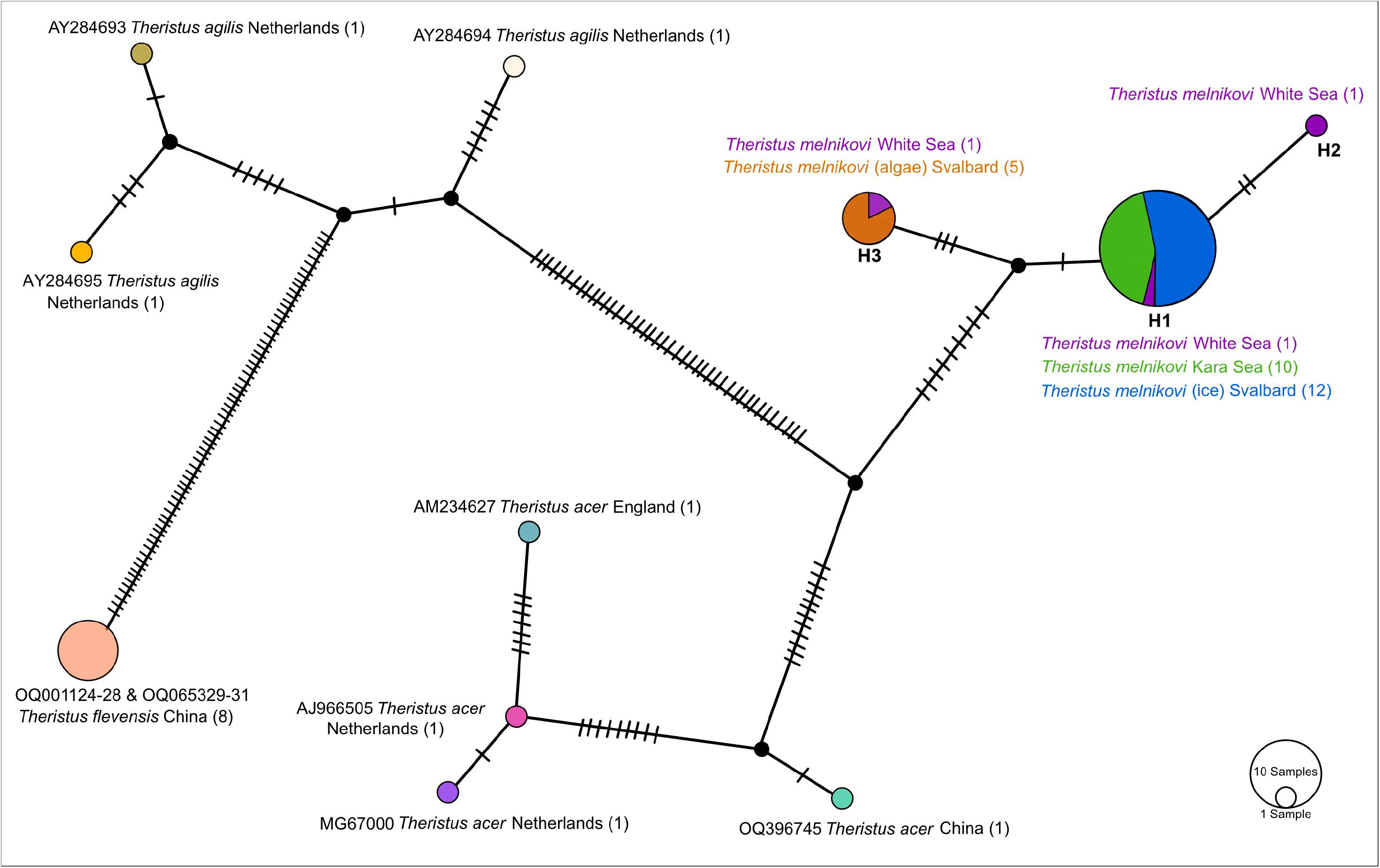
TCS haplotype network based on 18S rDNA sequences of *Theristus melnikovi* with close (*T. acer*) and distant relatives (*T. agilis* and *T. flevensis*). Each haplotype is coloured with the colour corresponding to the species and its sample location. The colours for *Theristus melnikovi* correspond to those of the Maximum Likelihood tree. The hash marks indicate the number of mutational steps between haplotypes. The size of the circle represents the sample size for each haplotype.

## Discussion

### Habitat-associated phenotypic variation in *Theristus melnikovi* revealed through integrative taxonomy

In this study, for the first time, we investigated the summer habitat of the sympagic nematode *Theristus melnikovi*, in the high Arctic fjords of Svalbard. Using an integrative taxonomic approach, we discovered that macroalgal-associated *Theristus* specimens morphologically resembling *Theristus acer* were genetically identical to sympagic *T. melnikovi.* Phylogenetic reconstruction of the complete 18S rRNA gene showed that specimens collected from macroalgae formed a monophyletic clade with *T. melnikovi* from sea-ice in Svalbard (this study; Pitusi et al., 2021), the Kara Sea and White Sea (Timchenko et al., 2024b) (Fig. 5). Additionally, sequenced juvenile macroalgal specimens shared an 18S haplotype with a singular male of *T. melnikovi* from the White Sea (19TMWS; ON153225) (Fig. 6). Together, these molecular data supported the assignment of the macroalgal specimens to *T. melnikovi* rather than *T. acer*, and we suggest that they represent a habitat-associated phenotypic variation of *T. melnikovi*.

We acknowledge that species assignment based on a single genetic marker should be interpreted cautiously, particularly because 18S rRNA is relatively conserved and insufficient to investigate mechanisms underlying morphological variation. We attempted to amplify the more variable 28S rRNA region but obtained only a single sequence from a sea-ice specimen, precluding the comparison between sea-ice and macroalgal-associated individuals. Nevertheless, the use of the full-length 18S rRNA rather than partial sequences, provided us with stronger evidence that the macroalgal specimens belong to the *T. melnikovi* lineage rather than *T. acer*.

Despite their genetic similarity, morphological examination of macroalgal specimens placed them in an intermediate position between ice-associated *T. melnikovi* and published records of the closely related benthic species *T. acer.* Compared with ice-associated *T. melnikovi*, the macroalgal-associated specimens had shorter, stouter bodies, proportionally longer tails, and smaller anterior structures (Fig. 4 A-D), resulting in an appearance more similar to that of *T. acer.* However, the morphological ranges of the Svalbard macroalgal specimens and published records only partially overlapped. For genital measurements, the overlap in both males and females was closest for landfast ice and macroalgal specimens. This illustrated the difficulty in species delimitation in closely related *Theristus* species based on body morphology alone. Additionally, cryptic speciation has previously been demonstrated for *T. acer* from the Netherlands (Derycke et al., 2013), and for *T. melnikovi* in the White Sea (Timchenko et al., 2024b), which emphasises the need for integrative taxonomy.

Morphological variation was also evident among sea-ice and macroalgal juvenile specimens, raising the question of when habitat-associated phenotypic differentiation arises during development. However, this study did not differentiate between juvenile developmental stages, so we could not determine at which stage differences emerged. We also did not identify the most useful morphological character to distinguish barcoded and mounted macroalgal specimens, as all sequences were from juveniles. Nevertheless, the occurrence of morphological differentiation in individuals that are genetically similar suggests that environmental conditions could contribute to the observed phenotypic differences.

Phenotypic plasticity is well-recognised in nematodes (e.g. Derycke et al., 2008), although it has received less attention in free-living marine taxa compared to free-living terrestrial or host-associated species (e.g. Kiontke & Fitch, 2010; Sommer et al., 2017). Reports of phenotypic plasticity in free-living nematodes are scarce, but a small number of studies have documented pronounced morphological variation despite low genetic variation (Miljutin & Miljutina, 2016; De Oliveira et al., 2017; Hauquier et al., 2017). De Oliveira et al. (2017), for example, described populations of *Paracanthonchus gynodiporata* Apolonio Silva De Oliveira, Decraemer, Moens, Dos Santos & Derycke, 2017 over a 1,000-km stretch from the Brazilian coast that displayed low genetic variation but substantial differences in morphometric characters, including buccal cavity width, the position of the amphidial fovea, and cephalic-setae length. Similar morphological variation has also been reported in the deep-sea nematode *Acantholaimus* Allgén, 1933 (Miljutin & Miljutina, 2016). Thus, morphological variation despite limited genetic differentiation is well-established, making habitat-associated phenotypic variation in *T. melnikovi* biologically plausible.

The observed variation between specimens may reflect phenotypic plasticity induced by differences between sea-ice and macroalgal habitats. These habitats differ substantially in their physical structure and stability. Although landfast ice undergoes a seasonal cycle of growth and melt, the lower 10 cm of the ice, where *T. melnikovi* resides, remains relatively stable during much of the season with respect to temperature, salinity, and food availability (e.g. Pitusi et al., 2023). Structurally, the sea-ice matrix consists of a network of microscopic brine channels, which are size-restrictive (Krembs et al., 2000) and, thus, appear to favour a long and slender phenotype that we, and other studies (e.g. Timchenko et al., 2024b), have observed.

In contrast, macroalgae habitats are less likely to be size-selective compared to the sea-ice matrix. Although macroalgae may be exposed to wave action and currents, sheltered microhabitats can occur near the holdfast (Moore, 1971), where the alga attached to the substrate, and among the fronds of species such as *Fucus* (Heip et al., 1985). Such lack of size constraints in these microhabitats may contribute to the stouter phenotype observed in macroalgal-associated specimens. Alternatively, environmental conditions such as temperature, experienced in the different sampled fjords, may contribute to the observed variation in body morphology (Armenteros & Ruiz-Abierno, 2015).

### Macroalgae as a potential summer refuge for sympagic nematodes

Projected losses of Arctic sea-ice raise important questions about the future of ice-associated fauna, of which *T. melnikovi* is just one representative. In the Central Arctic Ocean, summer sea-ice is projected to disappear within the next decade (Heuzé & Jahn, 2024), while Svalbard fjords are expected to be ice-covered, if at all, for fewer than two months per year (Urbański & Litwicka, 2022). Understanding how sympagic species persist during the ice-free season is therefore increasingly important.

Although *T. melnikovi* was previously regarded as an endemic sea-ice nematode (Tchesunov, 1986), its occurrence in coastal sea-ice has changed this view (Tchesunov & Portnova, 2005; Pitusi et al., 2021; Timchenko et al., 2024b). However, this raised questions about the species’ life history and summer habitat. A few individuals have been reported from benthic sediment, although in very low abundances (Timchenko et al., 2024b; Timchenko et al., 2025). Our findings suggest that macroalgae may provide an alternative (summer) habitat, or refuge, for this species. Rather than being restricted to only sea-ice, *T. melnikovi* appears to be able to occupy and exploit macroalgal habitats.

Macroalgae may also provide a food-rich habitat. Nematode associations with macroalgae are well-documented across numerous genera, including *Theristus* (e.g. Jensen, 1984; Derycke et al., 2007). The holdfast, stipe, and blade create a heterogeneous habitat that can support microbial biofilms, accumulate detrital material, and provide a refuge from predators (Heip et al., 1985). Although the feeding ecology of *T. melnikovi* remains poorly resolved, based on gut-content observations (Tchesunov & Riemann, 1995) and its unarmed buccal cavity, it is considered a non-selective deposit feeder (Wieser’s Type 1B; Wieser, 1953). It has been suggested that its diet consists of bacteria, protists, and dissolved organic matter (Tchesunov & Riemann, 1995; Kramer, 2011). Our genomic data revealed the presence of gammaproteobacteria *Oleispira* and *Colwellia*, which are well-known bacteria from Arctic sea ice (Thiele et al., 2022; Dilliplaine et al., 2025), and thus could be a food source for *T. melnikovi.* During ice-free periods, macroalgal-associated biofilms could provide food resources for *T. melnikovi*, although dietary analyses are needed to confirm these suspicions.

## Conclusion

Based on the integration of morphological and molecular analyses, we determined that the sympagic nematode *Theristus melnikovi* occupies macroalgal habitats in Svalbard fjords. Both phylogenetic and haplotype reconstruction placed macroalgal-associated specimens within the *T. melnikovi* clade. Morphologically, the macroalgal-associated specimens displayed shorter and stouter bodies, proportionally longer tails, and smaller anterior structures. Together these findings demonstrate habitat-associated phenotypic variation within *T. melnikovi* that is consistent with phenotypic plasticity. Although the capacity of macroalgae to compensate for future sea-ice loss remains unknown, the occurrence of *T. melnikovi* populations in macroalgal habitats indicates greater ecological flexibility than previously known.

To distinguish habitat-associated phenotypic variation from genetically structured population divergence will require broader geographical sampling, more variable genetic markers, and ideally common-garden experiments.

## Acknowledgements

The authors would like to thank Kyle Dilliplaine for his donation of his 2015 sea-ice samples. The authors would like to thank the UNIS logistics department for winter field support, UNIS AB-320 course [project nr. 411320] and the crew of *R/V Helmer Hanssen* for making summer sampling possible, and Stuart Thompson for field-and labwork related support. Thanks to Elaine Runge for her support in the field in 2021, and Sara V. Widera for picking the two nematodes for genome sequencing, in 2024. Johan Nylander (Department of Bioinformatics and Genetics, Swedish Museum of Natural History) is thanked for software programming support. The authors acknowledge the use of the Freiburg Galaxy server, funded by the German Federal Ministry of Education and Research BMFTR grant 031 A538A de.NBI-RBC and the Ministry of Science, Research and the Arts Baden-Württemberg (MWK) within the framework of LIBIS/de.NBI Freiburg. Fieldwork was done under the permits issued by the Governor of Svalbard (Sysselmesteren på Svalbard).

## Contributions

CRediT: **Vanessa Pitusi:** Conceptualization, Funding acquisition, Investigation, Methodology, Formal analysis, Writing – original draft, Writing – review & editing; **Janne E. Søreide:** Conceptualization, Funding acquisition, Investigation, Writing – review & editing; **Rebecca J. Duncan:** Funding acquisition, Investigation, Writing – review & editing; **Oleksandr Holovachov:** Conceptualization, Methodology, Formal analysis, Writing – original draft, Writing – review & editing.

## Data availability statement

The data that support the findings of this study are openly available from NCBI GenBank under the accession numbers PZ803337-PZ803338 and PZ805121-PZ805139. Raw data and assembled nuclear and mitochondrial genome assemblies are available from the European Nucleotide Archive under the accession number PRJEB124229. Raw reads under BioSample numbers SAMEA123379195/SAMEA123379196 and accession numbers ERR17745088/ERR17745089 respectively. The nuclear genome is available under accession number ERZ29893501 and at https://www.ebi.ac.uk/ena/browser/view/GCA_988240845.1. The mitochondrial genome is available under accession number ERZ30198781 and at https://www.ebi.ac.uk/ena/browser/view/GCA_988301505.1. The two COX1 sequences are available from the NCBI GenBank under the accession numbers QB081567 and QB081568 respectively.

## Supplementary materials

**Supplementary Data S1.** 18S alignment used for phylogenetic analysis and reconstruction.

**Supplementary Data S2.** 18S alignment used for haplotype network reconstruction.

**Supplementary Data S3.** Sequencing, assembly and analysis of nuclear and mitochondrial genomes from *Theristus melnikovi*.

**Table S1.**
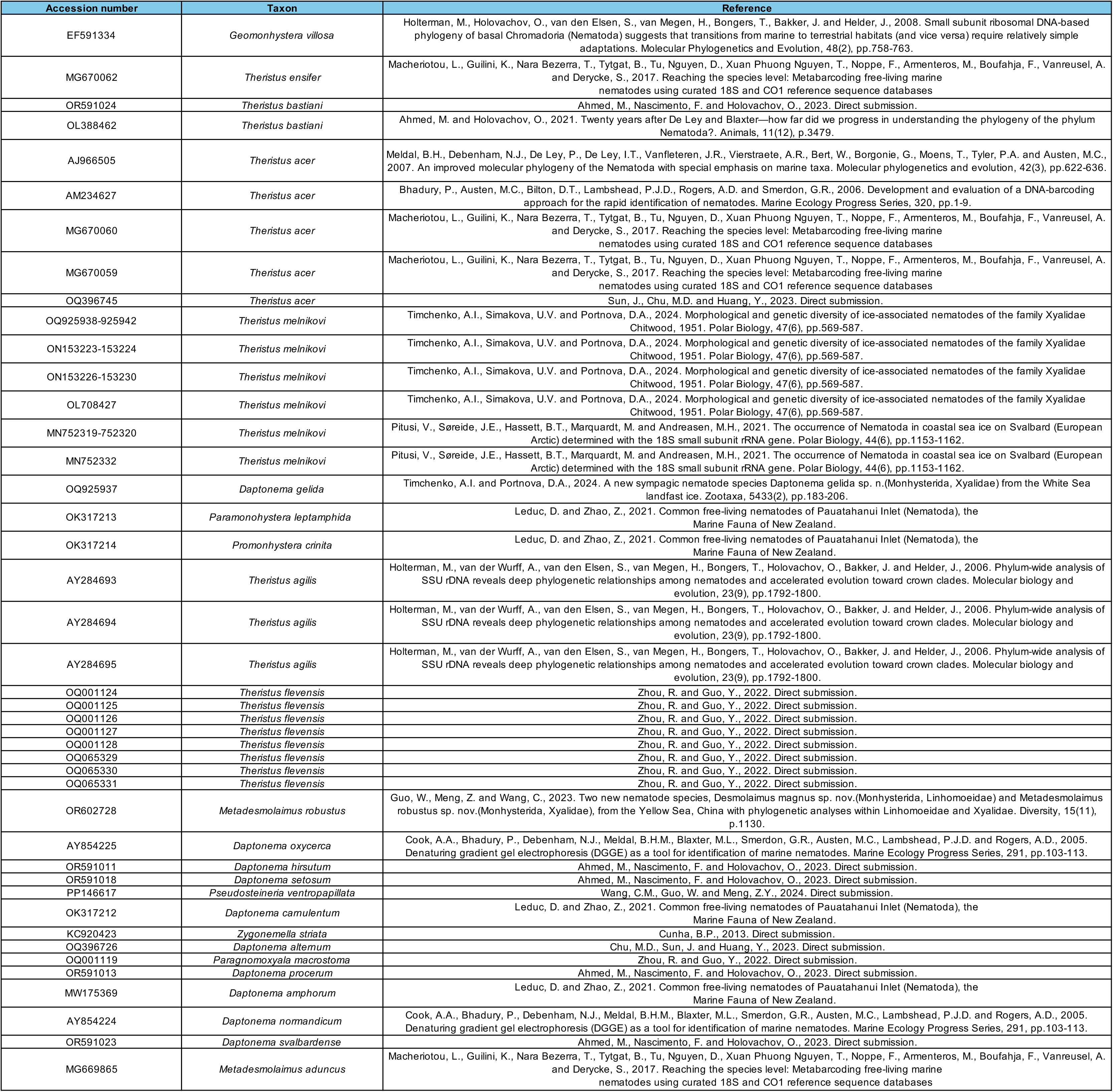
GenBank sequences downloaded and used in molecular analysis.

**Table S2.**
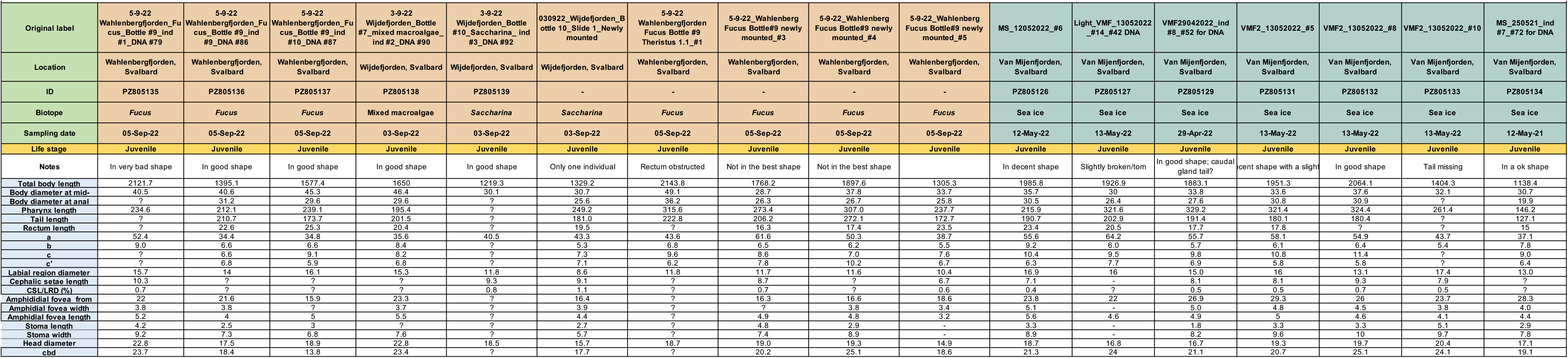
Morphometric measurements (in μm except the ratios a, b, c, c’, and where the percentage (%) is stated) of juvenile *Theristus* specimens from macroalgae and sea ice, in Svalbard. a = total body length/maximum body diameter, b = total body length/pharynx length, c = total body length/tail length, c’ = tail length/body diameter at anal, cbd = corresponding body diameter, - = absence of characteristic,= not possible to measure due to body deformation or incorrect positioning.

### Supplementary Data S3

#### Specimen collection

Nematodes were extracted from landfast sea ice samples collected in Van Mijenfjorden (77.7°N, 15.5°E) along the southwestern coast of Spitsbergen, in 2024. Sea ice samples were collected with a KOVACS ice corer (ø = 9 cm, Kovacs Ice Coring Equipment, USA). The lower 3-cm (ice -water interface) of each ice core was placed in a sampling bag and stored in a cooler while transported back to the lab. In the lab, 300 mL of 0.7-µm filtered seawater was added to 0-3 cm ice core section to prevent osmotic shock for the organisms inhabiting the ice core when melted in the dark at +4°C. The melted cores were concentrated over a 20-µm sieve and transferred to a petri dish. A Leica MZ16 stereomicroscope (Leica Microsystems, Germany) was used to locate and identify individual nematodes, which were picked with a glass Pasteur pipette and transferred to *RNAlater* solution (Thermo Fisher Scientific) in individual Eppendorf tubes. Specimens were subsequently identified using both Olympus SZX12 and Nikon Eclipse 80i microscopes. Morphological vouchers on permanent glass slides are deposited in the collection of the Arctic University Museum of Norway, Tromsø, Norway under accession numbers TSZY-548 – TSZY-553 (https://en.uit.no/tmu, contact person: V. Pitusi).

#### Multiple displacement amplification, sequencing and quality control

One female (GAI) and one male (GAS) of *Theristus melnikovi* were used to generate genomic data. Both specimens were collected from sea-ice in Van Mijenfjorden in 2024 as described above and preserved in *RNAlater* (Thermo Fisher Scientific). Subsequently, specimens were washed several times in DNAse/RNAse free water and placed in 4 µl of *REPLI-g Advanced sc Storage buffer* (Qiagen). Tissue digestion and whole genome amplification was performed separately for each individual nematode using *REPLI-g Advanced DNA Single Cell Kit* (Qiagen) following manufacturers protocol with modifications by Lee et al. (2024). After amplification, DNA was purified with *AMPure XP beads* (Beckman Coulter) and quantified using *Qubit™ dsDNA Quantification Assay Kit* (Thermo Fisher Scientific). 1.5 µg of the total amplified DNA was digested using *T7 Endonuclease* (Lee et al. (2024), purified with *AMPure XP beads* and quantified again using *Qubit™ dsDNA Quantification Assay Kit*. Sequencing was performed using ONT MinION, *R10.4.1 flow cell* (FLO-MIN114) and *Rapid Sequencing Kit V14* (SQK-RAD114) following manufacturers protocol. Post-run super accuracy (SUP) basecalling was done using latest version of *MinKNOW* (Oxford Nanopore Technologies). Palindromic chimaeras were identified and resolved using the *remove_palindrome.zsh* script (https://github.com/nylander/FASTQ_Palindrome_Cleaning_Pipeline) which is a modification of a script published by Lee et al. (2024). <u>usegalaxy.eu</u> server (The Galaxy community, 2026) was used for some of the subsequent steps. *fastplong* (Chen 2023) was used to remove sequencing adapters, trim 5’ and 3’ sequence ends and remove sequences with average phred quality less than 20 and shorter than 1000 bases. *GenomeScope* Galaxy ver. 2.1.0 (Ranallo-Benavidez et al., 2020) and *NanoPlot* ver. 1.46.2 (De Coster and Rademakers 2023) were used to assess the quality of the filtered raw reads.

#### Nuclear genome assembly and evaluation

*Flye* ver. 2.9.6 (Kolmogorov et al., 2020) installation on the usegalaxy.eu server was used to generate draft assemblies using both regular and metagenomic assembly modes, as well as varying the input quality of the reads (*--nano-raw*, *–nano-corr*, *–nano-hq*). Metagenome assembly mode with *--nano-hq* setting and three polishing iterations produced the most complete assemblies for both samples, which were mined for ribosomal genes using *barrnap* ver. 0.9 (https://github.com/tseemann/barrnap) to be used for haplotype network and phylogenetic analyses. For the final nuclear genome assembly, reads from both samples were combined. Draft assembly was generated using *Flye* ver. 2.9.6 (Kolmogorov et al., 2020) in metagenomic assembly mode using *--nano-raw setting*. *BlobToolKit* (Challis et al., 2020), was used to visualise, evaluate and decontaminate draft assembly: contigs identified as “chordata” and all non-metazoan contigs were removed (Figure S3-1). *BUSCO* ver. 6 (Tegenfeldt et al., 2025) was used to access the assembly completeness against four reference datasets (*nematoda_odb12*, *nematoda_odb12.1*, *chromadorea_odb12*, *chromadorea_odb12.1*). Annotation of protein coding genes was not performed due to absence of RNAseq data.

**Figure S3-1.**
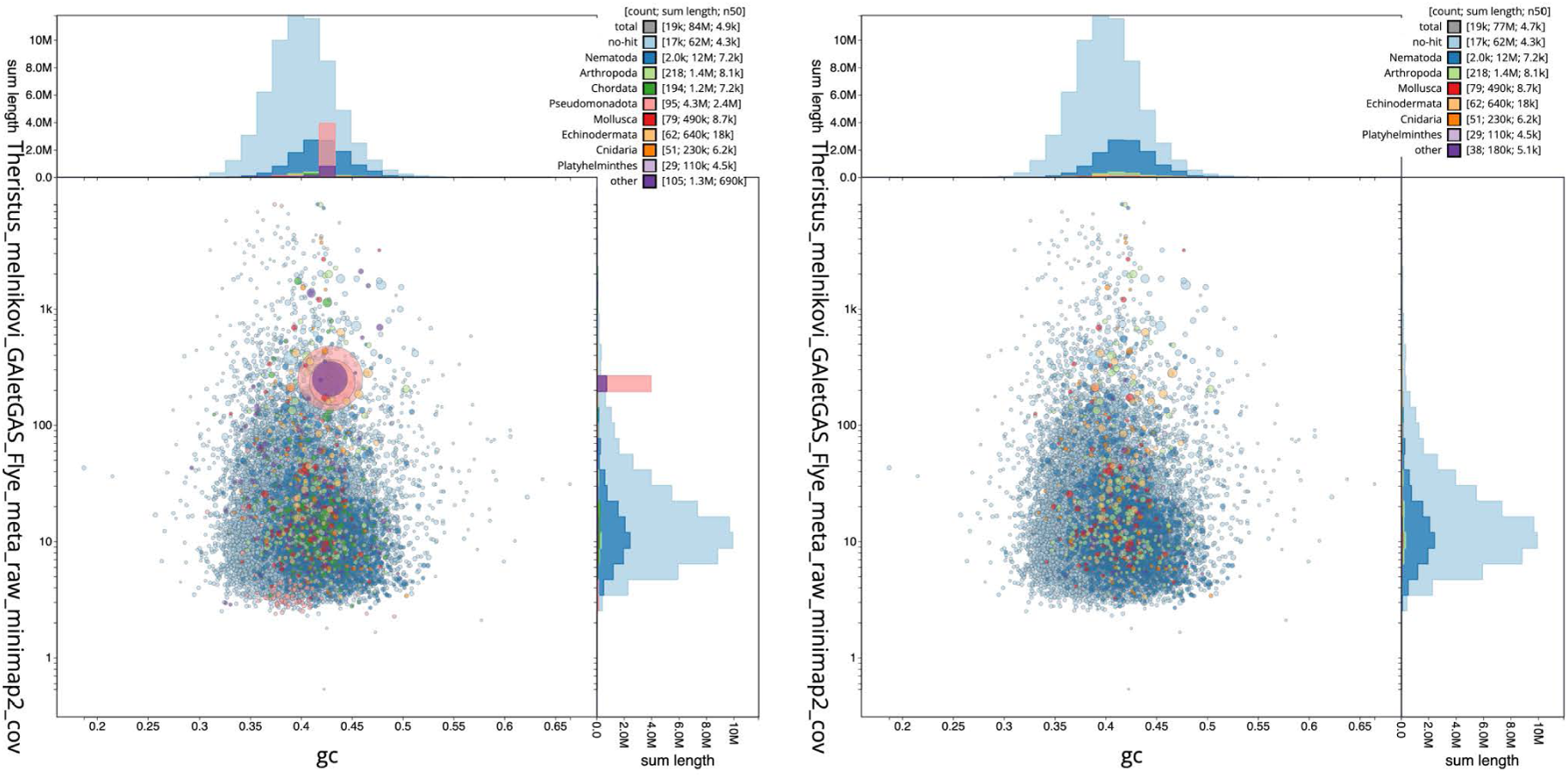
The *BlobToolKit* blob plot visualising sequence coverage (vertical axis) and GC content (horizontal axis) with the circles representing scaffolds, with the size proportional to scaffold length and the colour representing phylum while the histograms along the axes display the total length of sequences distributed across different levels of coverage and GC content, for original (left) and decontaminated (right) assemblies.

The final decontaminated genome assembly (Figure S3-2, Table S3-1) is only partial and very fragmented, having a total length of 77 Mb, consisting of 19027 contigs, with an N50 = 5 kb, with the longest contig being 92.4 kb. *BUSCO* ver. 6 (Tegenfeldt et al., 2025) analysis using various reference datasets identified only 18.9-29.5% of the expected genes.

**Figure S3-2.**
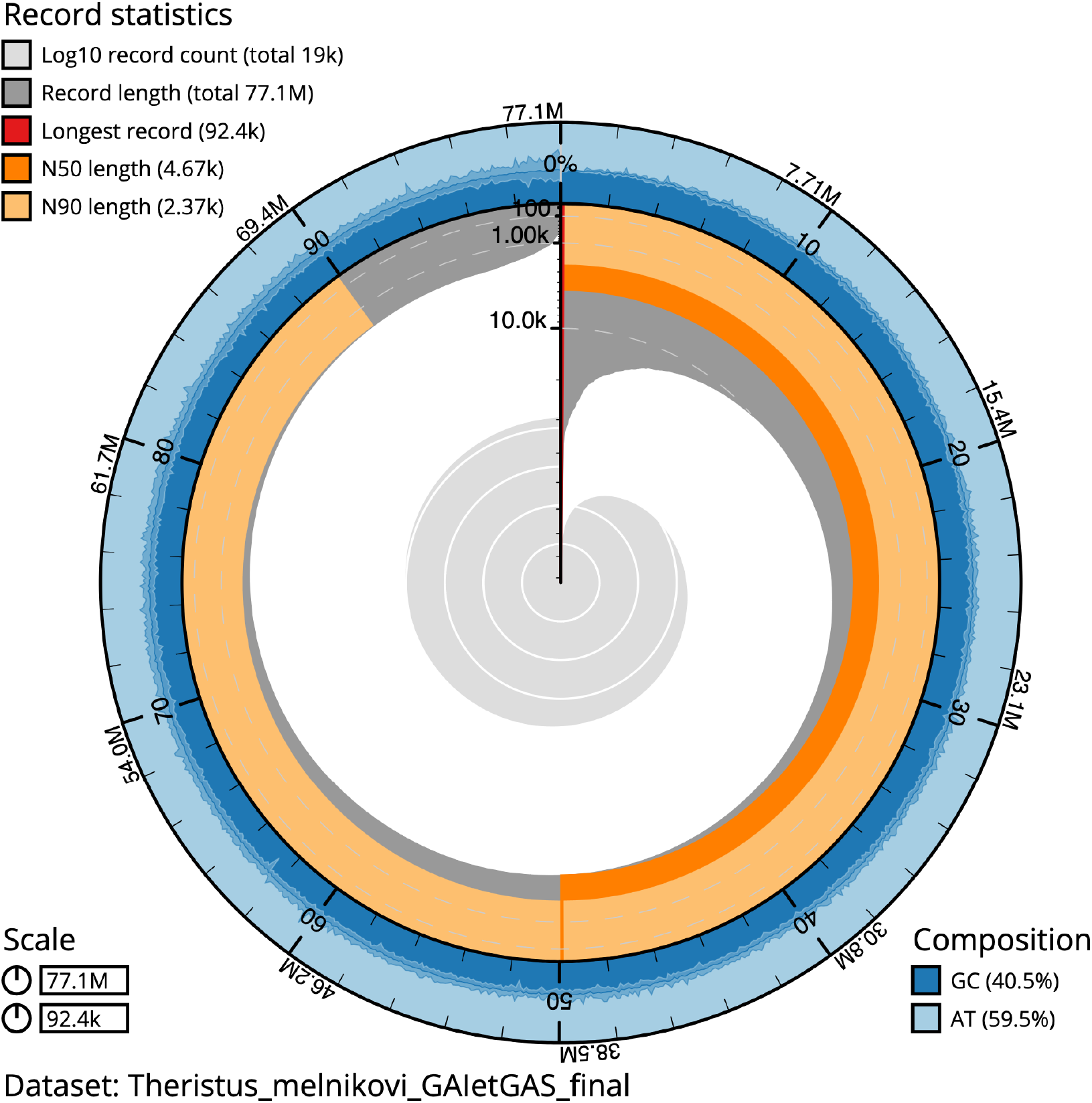
The *BlobToolKit* snail plot visualising the assembly metrics, the circumference represents the length of the whole genome sequence, the outermost blue tracks display the distribution of GC, AT, and N percentages, scaffolds are arranged clockwise from longest to shortest and are depicted in dark grey, the longest scaffold is indicated by the red arc, and the deeper orange and pale orange arcs represent the N50 and N90 lengths.

**Table S3-1.** Nuclear genome (draft) assembly statistics

| Table S3-1. Nuclear genome (draft) assembly statistics |  |  |  |  |
| --- | --- | --- | --- | --- |
| Accession | PRJEB124229 |  |  |  |
| Assembly | GCA_988240845.1 |  |  |  |
| Assembly level | contig |  |  |  |
| Span | 77 Mb (77081132 bp) |  |  |  |
| Number of contigs | 19027 |  |  |  |
| Contig N50 | 5 kbp |  |  |  |
| BUSCO: | nematoda<br>odb12<br>(596) | nematoda<br>odb12.1<br>(1477) | chromadorea<br>odb12<br>(1126) | chromadorea<br>odb12.1<br>(2034) |
| Complete (all) | 176 (29.5%) | 317 (21.5%) | 285 (25.3%) | 385 (18.9%) |
| Complete (single-copy) | 170 (28.5%) | 307 (20.8%) | 276 (24.5%) | 375 (18.4%) |
| Complete (duplicated) | 6 (1.0%) | 10 (0.7%) | 9 (0.8%) | 10 (0.5%) |
| Fragmented | 155 (26.0%) | 307 (20.8%) | 222 (19.7%) | 358 (17.6%) |
| Missing | 265 (44.5%) | 853 (57.8%) | 619 (55.0%) | 1291 (63.5%) |

High fragmentation of the final assembly can be explained by sub-optimal DNA preservation in *RNAlater* supplemented by unavoidable fragmentation of DNA during *T7 endonuclease digestion* (part of the Multiple Displacement Amplification protocol) and tagmentation during library preparation (part of the *Rapid Sequencing Kit V14* protocol).

#### Mitochondrial genome assembly, annotation and evaluation

Mitochondrial genomes were assembled for each sequenced specimen separately following the strategy described in Morffe et al. (2026) and using <u>usegalaxy.eu</u> server (The Galaxy community 2026) for majority of the steps. Nucleotide sequences of nematode mitochondrial genes were downloaded from the https://github.com/WormsEtAl/Nematode-Mitochondrial-Database (Gendron et al. 2023) and converted into a *BLAST*-compatible nucleotide mitochondrial reference database with the <u>usegalaxy.eu</u> installation of a NCBI BLAST+ *makeblastdb* (Altschul et al. 1990). Amino acid sequences of protein-coding genes were extracted from the mitochondrial genomes of 52 nematode species across the entire phylum downloaded from the NCBI GenBank, and converted into a *Diamond*-compatible amino-acid mitochondrial reference database with the <u>usegalaxy.eu</u> installation of *Diamond makedb* (Buchfink et al. 2021). *blastn* (Altschul et al. 1990) with *E-value* cutoff 0.001 was used to search the filtered raw reads against the nucleotide mitochondrial reference database, while *Diamond blastx* (Buchfink et al. 2021) set to invertebrate mitochondrial code and *E-value* cutoff 0.001 was used to search the filtered raw reads against the aminoacid mitochondrial reference database. *BBedit* was used to remove duplicate hits, and create a list of read sequence headers for the *seqkit* ver. 2.8.0 (Shen et al. 2024), which was used to pull relevant reads from the filtered raw reads file. Three input datasets, one based on *blastn* hits, one based on *diamond blastx* hits and one using combined hits were used as an input for the mitogenome assembly with *Canu* ver. 2.3 (Koren et al. 2017). Multiple assemblies were generated using the following constant settings: *technology* = Nanopore; *maximum corrected overlap mismatch* = 0.2; *target coverage for corrected reads* = 10; *low coverage depth* = 5; estimated genome size = 25k. Five combinations of *minimum read length* / *minimum overlap* were used in parallel: 1000/250, 1000/500, 2000/1000, 3000/1500 and 4000/2000. *Mitos2* (Bernt et al. 2013) was used to extract *COX1* from several randomly selected assemblies with the following settings: *genetic code* = Invertebrate (5), *reference data* = refseq63m. This *COX1* sequence was then identified using NCBI *blastn* against GenBank core nucleotide database (excluding uncultured and environmental sample sequences). Same *COX1* sequence was used to positively identify remaining assembled contigs as belonging to the same target species, orient them in the same direction, align and circularise using *AliView* ver. 1.30 (Larsson 2014). Draft consensus assembly was created by combining all aligned mitochondrial contigs using majority rule approach. Only one of the assemblies was circularised and was analysed further. The second assembly was partial, but aligned perfectly with the circular assembly except for the gap. *minimap2* (Li 2021) was used to map raw reads against draft assembly using *map-ont* setting while *pilon* ver. 1.20 (Walker et al. 2014) was used to improve it based on *minimap2* output. *Tandem repeats finder* (Benson 1999) and Claude AI by Anthropic were used to identify repetitive regions. *Mitos2* (Bernt et al. 2013) and *Aragorn* ver. 1.2 (Laslett and Canback 2004) were used to annotate final mitochondrial genome assembly. *samtools* (Danecek et al. 2021) was used to estimate the read coverage from the *minimap2* output. *Proksee* (Grant et al. 2023) was used to visualise the final assembly, further edited in *Affinity Studio 2* (Figure S3-3).

**Figure S3-3.**
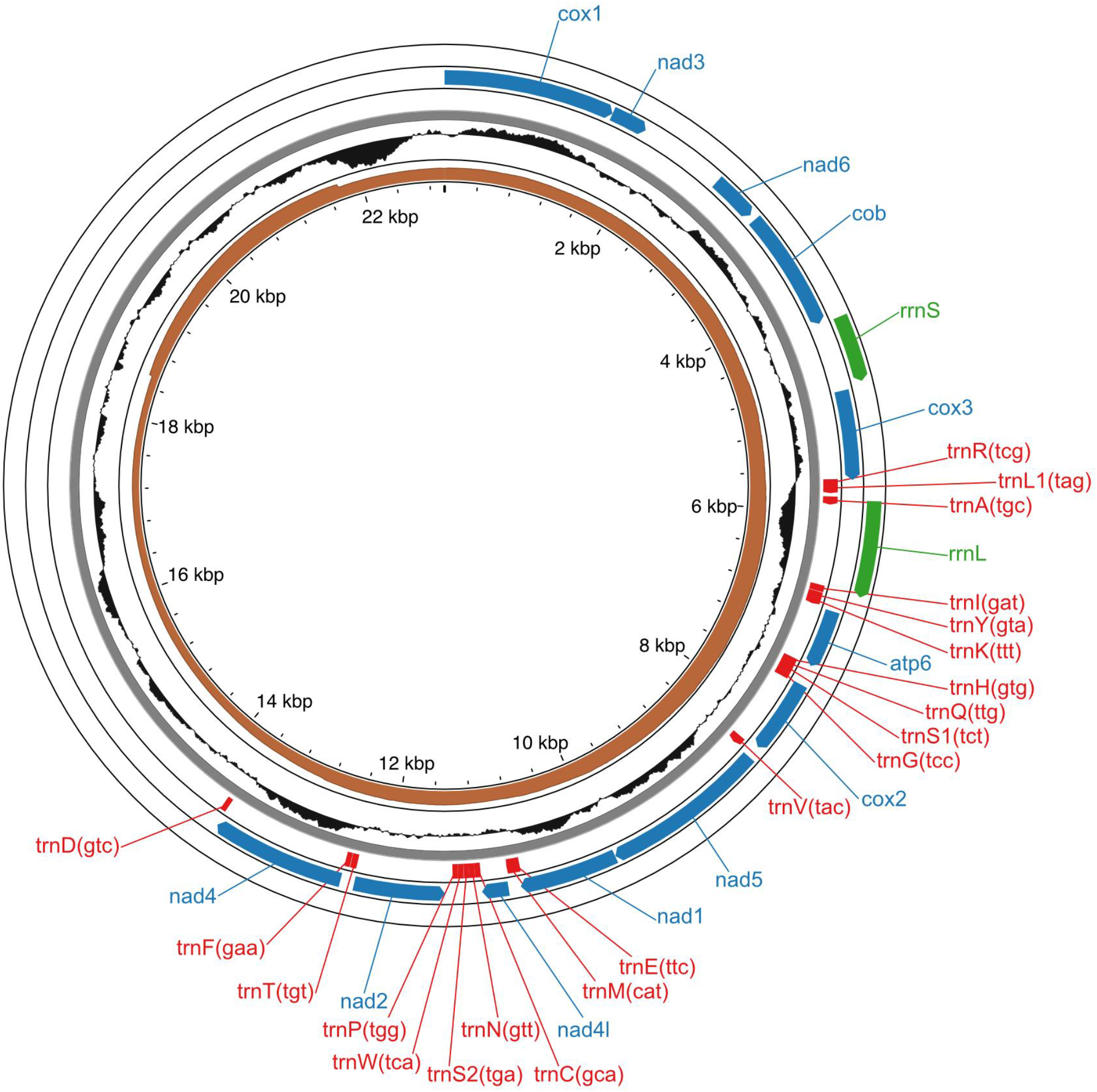
Annotated mitochondrial genome of *Theristus melnikovi* Tchesunov, 1986 showing protein coding genes (CDS, blue; n=12), ribosomal RNA genes (green, n=2), transfer RNA genes (red; n=18), read coverage (brown) and GC content (black).

The complete mitochondrial genome has a total length of 22,976 bases (Figure S3-3). Mitogenome assembly has two separate compact tandem-repeat arrays: first block ∼1.85 kb long occupying 13,818– 15,667 bp, composed of short ∼209 bp unit tandemly repeated 8–9 times; second block ∼4.4 kb long, occupying 18,568–22,976 bp, composed of ∼860–970 bp unit tandemly repeated five times. Annotation using *Mitos2* (Bernt et al. 2013) and *Aragorn* ver. 1.2 (Laslett et al. 2004) identified a complete set of 12 protein coding genes (*atp6*, *cob*, *cox1-3*, *nad1-6*, and *nad4L*), two rRNA genes and 18 tRNA genes (Figure 2). Single tRNA gene, *tRNA-Leu (L2)*, could not be annotated. We do not postulate that this tRNA is completely missing from the mitogenome of *T. melnikovi*, but that it can be annotated in the future, when more mitogenomes for species of the family Xyalidae are available.

#### *COX1*-based phylogeny

The maximum likelihood *COX1*-based phylogeny was generated as follows: partial *COX1* sequences from species of the family Xyalidae were combined with *T. melnikovi* sequences (68 partial *COX1* sequences from at least 37 species, many of which are not identified), aligned and trimmed using *AliView* ver. 1.2 (Larsson 2014); *IQ-TREE 2.3.1* macOS-arm version (Minh et al. 2020) was used to infer the tree using *ModeFinder* (Kalyaanamoorthy et al. 2017) to estimate the best mitochondria-specific substitution model, followed by standard nonparametric bootstrap (1000 replicates). Cladogram was edited in *TreeViewer* ver. 2.2 (Bianchini and Sánchez-Baracaldo 2024) and *Affinity Studio 2* (Figure S3-4).

**Figure S3-4.**
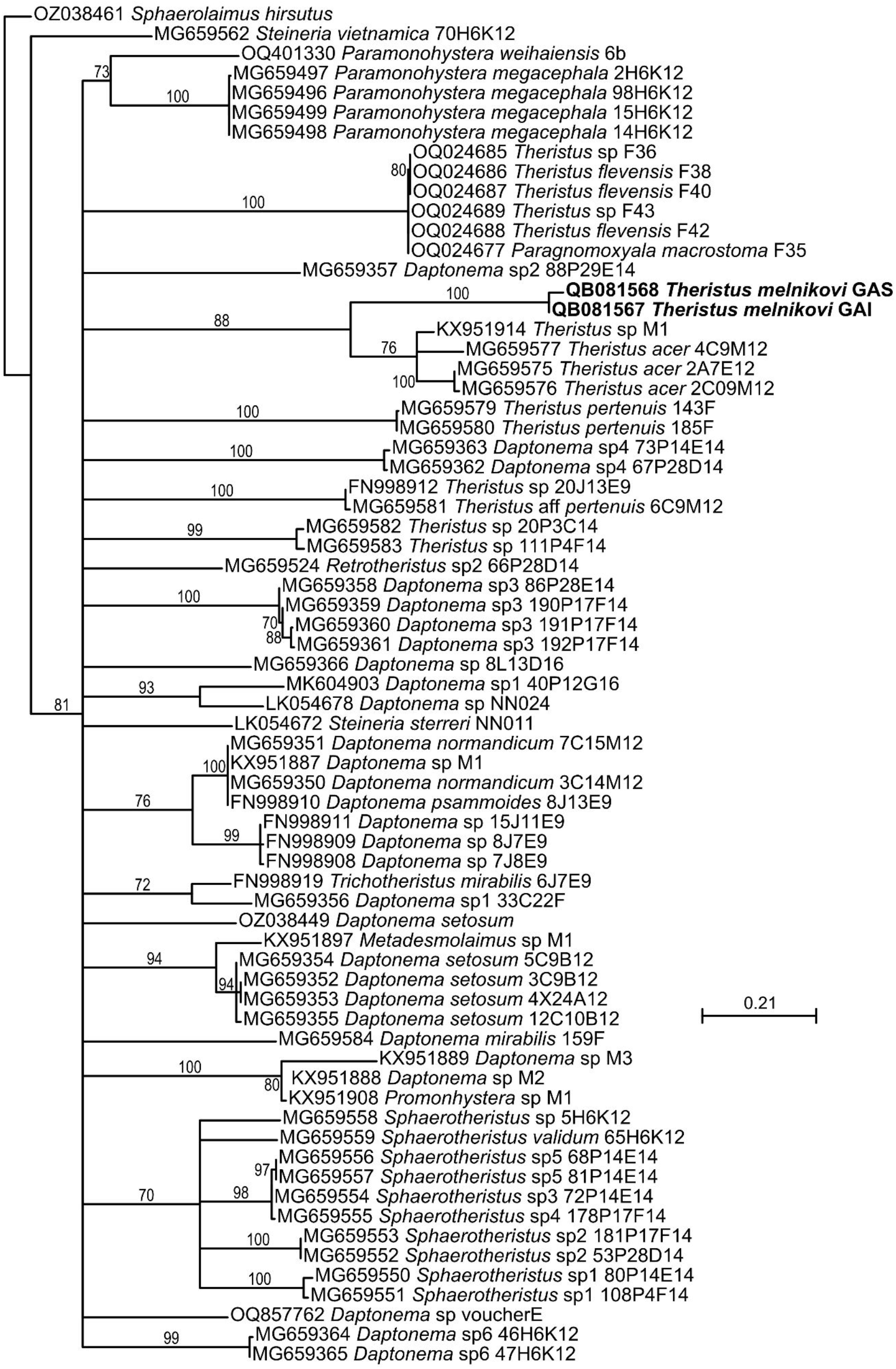
Maximum likelihood (ML) tree inferred from the partial *COX1* gene for species from the family Xyalidae, including *Theristus melnikovi* Tchesunov, 1986 from this study, with *Sphaerolaimus hirsutus* Bastian, 1865 (Sphaerolaimidae) used as an outgroup. All three sequences of *Theristus acer* in the tree originate from specimens collected in the Netherlands, *Theristus* sp. M1 was collected along the Atlantic coast of Portugal.

Maximum likelihood phylogeny (Figure S3-4) is poorly resolved, but places *T. melnikovi* and *Theristus acer* Bastian, 1865 as sister taxa in a strongly supported clade, supporting close relationships between the endemic Arctic sea ice species (*T. melnikovi*) and the species widely distributed in a range of marine habitats in the northern hemisphere (*T. acer*).

